# Characterization of cell-fate decision landscapes by estimating transcription factor dynamics

**DOI:** 10.1101/2022.04.01.486696

**Authors:** Sara Jiménez, Valérie Schreiber, Reuben Mercier, Gérard Gradwohl, Nacho Molina

## Abstract

Modulation of gene expression during differentiation by transcription factors promotes cell diversity. Despite their role in cell fate decisions, no experimental assays estimate their regulatory activity in a high-throughput manner and at the single-cell resolution. We present FateCompass for identifying lineage-specific transcription factors across differentiation. It uses single-cell transcriptomics data to infer differentiation trajectories and transcription factor activities. We combined a probabilistic framework with RNA velocities or a differentiation potential to estimate transition probabilities and perform stochastic simulations. Also, we learned transcription factor activities using a linear model of gene regulation. Considering dynamic changes and correlations, we identified lineage-specific regulators. We applied FateCompass to an islet cell formation dataset from the mouse embryo, and we found known and novel potential cell-type drivers. Also, when applied to a differentiation protocol dataset towards beta-like cells, we pinpointed undescribed regulators of an off-target population, which were experimentally validated. Thus, as a framework for identifying lineage-specific transcription factors, FateCompass could have implications on hypothesis generation to increase the understanding of the gene regulatory networks driving cell fate choices.

**Highlights:**

- We developed FateCompass, a flexible pipeline to estimate transcription factor activities during cell-fate decision using single-cell RNA seq data.
- FateCompass outlines gene expression stochastic trajectories by infusing the direction of differentiation using RNA velocity or a differentiation potential when RNA velocity fails.
- Transcription factor dynamics allow the identification of time-specific regulatory interactions.
- FateCompass predictions revealed known and novel cell-subtype-specific regulators of mouse pancreatic islet cell development.
- Differential motif analysis predicts lineage-specific regulators of stem cell-derived human β- cells and sheds light on the cellular heterogeneity of β-cell differentiation protocols.
- Experimental validation supports the proposed GRN controlling SC-EC differentiation predicted by FateCompass.

## Introduction

Gene regulation is pivotal during many biological processes, including development, cell cycle, regeneration, reprogramming, and cancer, and it usually occurs in a cell- and stage-dependent mode ^1^. Notably, cells transition from a less to a more differentiated state during differentiation via the interplay of transcriptional regulation events in a highly dynamic manner ^1,2^. Transcription factors (TFs) are essential proteins that have the ability to bind specific DNA regulatory regions and link signaling transduction networks to gene-specific transcriptional regulation ^3^; hence they are commonly used as readouts of pathway activities. Currently, there are no high-throughput techniques to measure TF activity; instead, their direct product, gene expression level, can be measured with an unprecedented high resolution using single-cell transcriptomics.

Single-cell RNA sequencing (scRNAseq) techniques allow identifying different cell types and, more importantly, the study of lineage-specification at the single-cell resolution ^4^. The inherent asynchrony of scRNAseq data has allowed the development of several approaches to reconstruct differentiation trajectories, which rely on variation among cell types within the captured population ^5^. The developed computational techniques include pseudotime methods ^6,7^ and RNA velocity ^8,9^. Noteworthy, pseudotime algorithms depend on the previous knowledge of the initial state, and it is limited to the analysis of general trends of biological progressions rather than the precise dynamics of individual cells. Conversely, RNA velocity overcomes the limitation of the directionality by leveraging the splicing kinetics and predicting the RNA expression states in the near future. Nevertheless, it has intrinsic limitations, e.g., when the spliced-to-unspliced mRNA ratio is trendless or in predicting the continuous evolution of cells over a long period of time ^10,11^. Typically, differentiation trajectories are used together with differential gene expression analysis to identify TFs specific to a given cell type ^12^. However, this approach ignores the fact that even lowly expressed TFs can have high regulatory activity, and it does not consider direct regulatory interactions with target genes. On the other hand, several methods to derive mechanistic signatures in cell-fate decisions from transcriptomics data have been proposed, including ISMARA ^13^ and DOROTHEA ^2^ for bulk RNA seq; and SCENIC ^14^ and metaVIPER ^15^ for scRNAseq. However, except for ISMARA, they are based on correlations between the expression of TF transcripts and the TF target genes or the expressed genes in general. Using correlations requires further assumptions or perturbation assays to distinguish causal relationships. Furthermore, none describes the dynamic change of TF activity throughout the cell-fate decision process, which is pivotal in time-dependent systems. The challenge is to devise a robust workflow that infers cell-type-specific regulators dynamically.

Here, we present FateCompass, a workflow that aims to identify lineage-specific TFs for a system undergoing differentiation. First, we outlined differentiation trajectories from progenitor cells to final states using a discrete Markov Process on a network. This allows us to describe stochastic gene expression dynamics during the cell fate decision process incorporating RNA velocity or differentiation potentials to infuse the differentiation direction. Then, we inferred TF activities by modeling the observed gene expression as a linear combination of the regulatory sites and the TF activity. Finally, we performed a differential TF activity analysis using statistical criteria. We applied FateCompass to a pancreatic endocrine differentiation system, where endocrine progenitors, marked by the transient expression of the TF Neurog3, differentiate towards glucagon-producing α-cells and insulin-producing β-cells, among others ^16^. We analyzed a well-characterized scRNAseq dataset from the developing mouse pancreas ^17^. FateCompass identified known and novel lineage-specific regulators; among them, Arx and Nkx6-1 as α- and β-specific, respectively. Further, to demonstrate the capabilities of FateCompass, we used a scRNAseq experiment from the differentiation of human stem cells towards pancreatic β-like cells ^18^. Of note, this complex population includes, besides the expected endocrine cells, an off-target population of intestinal cells called enterochromaffin (EC). FateCompass identified not only α- and β-specific but also EC-specific known and novel factors such as CDX2, for which we present further experimental support. FateCompass boosts the ability to identify lineage-specific regulators by estimating TF activities dynamically, revealing time-specific transcriptional regulatory interactions underlying cell-subtype specification.

## Results

### Inferring dynamic transcription factor activity during cell subtype specification

The FateCompass workflow aims to identify lineage-specific transcription factors for a cellular system undergoing differentiation. To mechanistically understand the dynamic transcriptional interactions underlying a cell subtype specification process, we designed a three-step pipeline using scRNAseq (Fig. 1). First, to depict the system’s dynamics, we outlined differentiation trajectories from progenitor cells to final states using a discrete Markov Process on a network representing cell states and possible transitions. Next, we focused on TFs as readouts of pathway activity due to their direct role in cell-specific transcriptional regulation. Hence, we inferred TF activities using a linear model of gene regulation, as in the original framework, ISMARA ^13^; we modeled the observed gene expression as a linear combination of the regulatory sites and the motif activity. Finally, to coarse-grain the list of TFs and identify lineage-specific regulators, we defined a differential motif activity analysis using three metrics.

**Figure 1.**
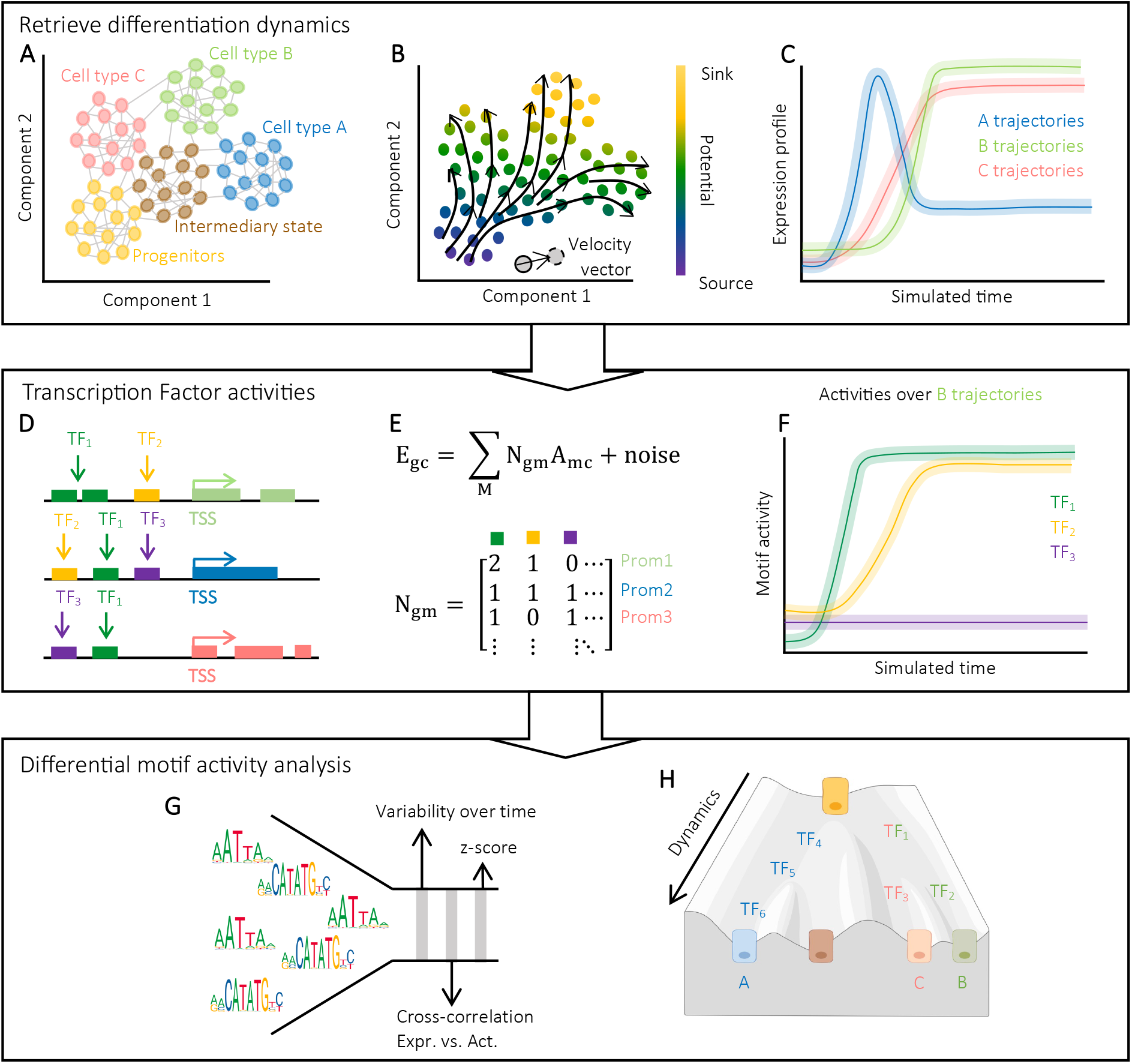
Workflow to decipher key transcriptional regulators underlying cell-fate decision landscape. **A**. scRNAseq data annotated for different cell-types and embedded in a low- dimensional space showing connections to the *k* nearest neighbors, which are computed on the first D dimensions (D<*k*) of the low-dimensional embedding. **B**. The direction of difierentiation, either from RNA velocity or Potential energy, is the underlying force to compute transition probabilities between cell states. **C**. The dynamic profile of a given gene over the trajectories ending in a respective final state (A, B, and C). We used stochastic simulations with the previously computed transition probabilities to infer the dynamic profiles. **D**. Computationally predicted transcription factor binding sites for difierent motifs across promoters. **E**. A linear model of gene regulation where the observed gene expression (E_gc_) is modeled as a linear combination of the regulatory sites (N_gm_) and the TF activity (A_mc_). **F**. The dynamic profile of the motif activity for TF_1_, TF_2_, and TF_3_ over the stochastic trajectories ending in the final state B. **G**. Difierential motif activity analysis. We selected the key TFs for each final fate according to three criteria: high variability over time, a high dynamic correlation between the motif activity and the expression of its mRNA, and a high z-score. **H**. We identified TFs that are important for defining several fates and TFs that are lineage-specific, and we order them according to who is active first over the dynamic profiles.

To infer the dynamic trends of the system undergoing differentiation and describe stochastic gene expression profiles along the cell-fate decision process, we used cell-to-cell similarity based on transcriptomic profiles together with RNA velocity ^8^ or a differentiation potential from progenitor to mature cells ^19^. Of note, single-cell transcriptomics provides a snapshot of intrinsically asynchronous cells that can be ordered based on similarities in the expression patterns to capture the time-evolution faithfully over the transition populations ^4,20^. Indeed, most trajectory inference methods using scRNAseq data assume that a cell change states in small transcriptional steps ^5-7,21^. Similarly, FateCompass uses this assumption to model differentiation trajectories, except that we implemented a cell-dependent drift that biases the trajectories towards the direction of differentiation. Briefly, we represented the phenotypic manifold in a low-dimensional space using a Markov chain on a network (Fig 1A and Methods 1.1.1). To that end, we first embedded the gene expression data in a significant low-dimensional manifold using Uniform Manifold Approximation and Projection (UMAP) ^22^. Notably, UMAP was first developed for clustering and has proven to be more performant than other non-linear dimensionality reduction algorithms when the embedding dimension is higher than two ^22,23^. Next, we built a nearest-neighborhood graph in the low-dimensional space connecting each cell with the n most similar neighbors. Currently, RNA velocity is a well-accepted method to infer differentiation dynamics unbiasedly; however, some of the method’s limitations lead to inconclusive velocity fields in some biological systems. For instance, some datasets might have, to name just a few, time frames out of the initial modeling framework, insufficient unspliced counts in the key biological driver genes, and multiple kinetic regimes ^10^. Therefore, to make FateCompass flexible and applicable to any differentiating system, we used either the RNA velocity vector (Methods 1.1.3) or the gradient of Potential energy from progenitor cells to mature cells (Methods 1.1.4) to bias the transitions between states in the Markov process (Fig 1B). The resulting transition probabilities reflect both transcriptional and directional similarities. To ultimately describe the time evolution of the differentiating system, we used a Monte Carlo sampling algorithm where the next-jump probability was given by the transition matrix of the Markov chain (Methods 1.1.5). This approach is instrumental in estimating quantities of interest, e.g., gene expression or transcription factor activity over differentiation trajectories (Fig 1C&F and Methods 1.1.6).

To decipher the transcriptional interactions driving cell subtype specification, we used TFs as a proxy because of their direct role in gene-specific transcriptional regulation ^24^. To predict TF activities during differentiation, we reasoned that changes in the transcriptional state, in response to developmental cues, are conditioned by conserved regulatory mechanisms – such as the interaction between TFs and promoters (Fig 1D). Similar to Balwierz et al. (2014), we used a linear model to infer TF activities (Methods 1.2.2). The primary assumption is that the transcription rate is controlled by the TF binding sites present in the promoters (Fig 1E and Methods 1.2.1); we considered promoters because of their direct assignation to the target genes based on their proximity to the transcriptional start site (TSS). There is no standard high-throughput way to assign reliably long-distant regulatory interactions such as enhancers to target genes. We defined a promoter region as the 2 kb region centered in the TSS. Importantly, we implemented a new regularization technique using data diffusion to control the model’s complexity and avoid overfitting (Methods 1.2.3). Shortly, we used the *k-*nearest neighborhood graph to smooth the learned activities correcting for dropout and other noise sources ^25^. In the data diffusion regularization, the cells share information through the local neighbors by a process analogous to diffuse the data over the network. The *t*-step is akin to raising the diffusion operator to the *t*^*th*^ power. We fitted the optimal value of *t* using a cross-validation scheme.

Finally, to identify lineage-specific TFs, we defined a differential motif activity analysis based on three metrics (Fig 1G and Methods 1.3). Firstly, we reasoned that activities that are important to explain the expression variation across cells should be relevant in the cell-fate decision; we summarize this using the z-score (Methods 1.3.1). Next, we detected TFs whose activity profile over the differentiation trajectories was highly changing-intuiting that these will have a crucial role in the state-transition process (Methods 1.3.2). Lastly, we acknowledge that for a TF to be active, it has first to be expressed; hence, we looked for TFs with high and positive dynamical cross-correlation (Methods 1.3.3). FateCompass boosts the ability to identify lineage-specific regulators by estimating TF activities dynamically. The potential to identify relative-time-specific regulatory interactions opens the door for hypothesis generation on transcriptional regulation cascades underlying cell-subtype specification processes (Fig 1H).

### Delineating transcriptional regulators during mouse islet cell formation

To assess the robustness of our workflow, we applied it to a well-characterized scRNAseq data set from the developing mouse pancreas with transcriptome profiles sampled from embryos at 15.5 days post-coïtum (E15.5) ^17^. In the Pancreas, endocrine cells differentiate from endocrine progenitors marked by the transient expression of the basic helix loop helix (bHLH) transcription factor Neurog3 ^16^. Bastidas-Ponce et al. (2019) profiled pancreatic epithelial cells using a Neurog3-Venus fusion reporter mouse line, sequencing both Venus-positive and Venus-negative (Epcam+) cells using droplet-based scRNAseq (10X genomics chromium). We decided to test the capabilities of our workflow with the data from E15.5 (3696 cells) because at this time point, endocrine cell commitment ends in four major cell types: glucagon-producing α-cells, insulin-producing β-cells, somatostatin-producing δ-cells, and ghrelin-producing ε-cells, Fig. 2A. Moreover, this dataset presents a strong directional velocity flow towards the final endocrine fates, Fig. 2A ^9^.

**Figure 2.**
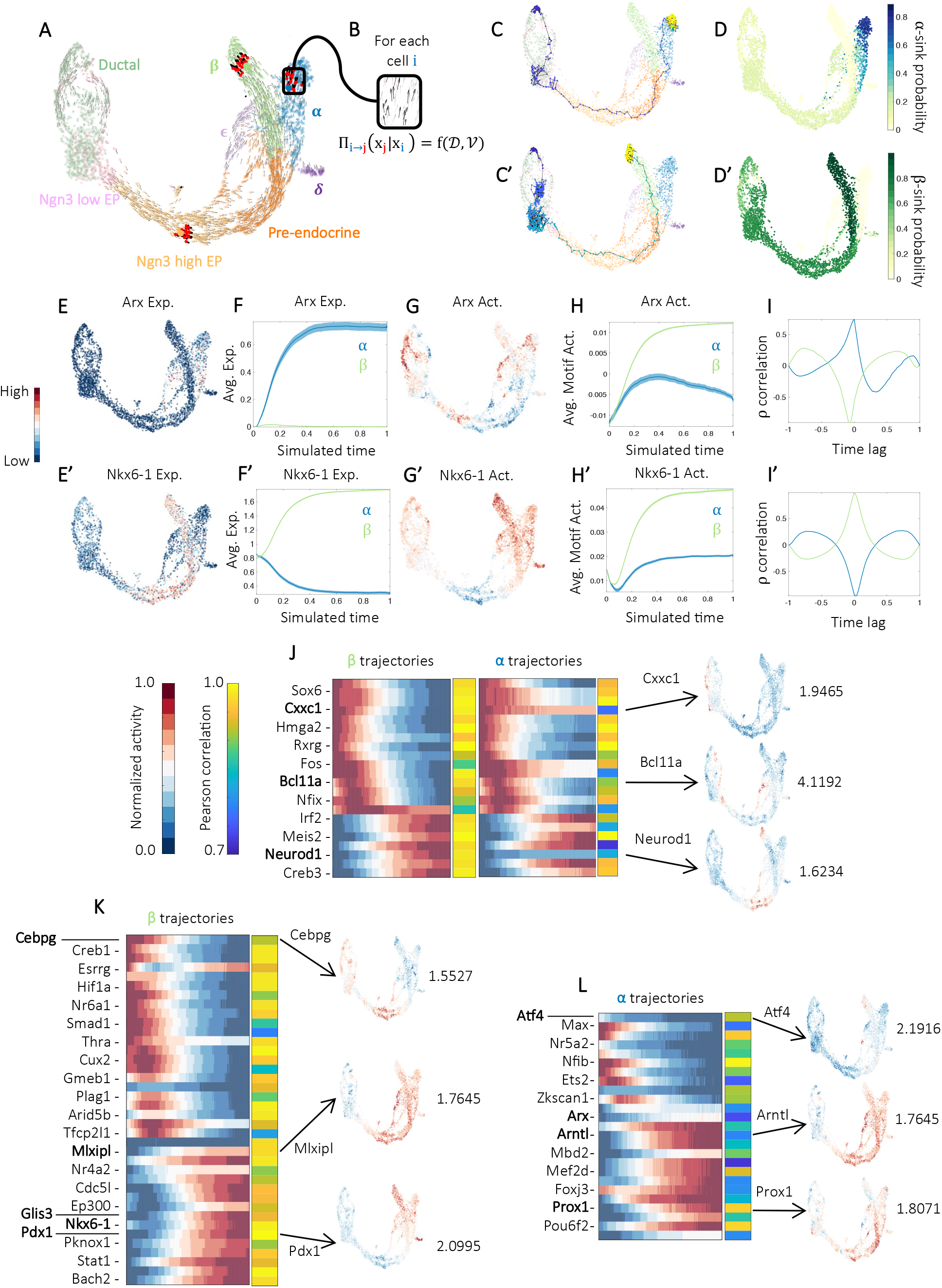
Islet cell formation landscape in the mouse. **A**. UMAP plot of 3696 cells at E15.5 from Bastidas-Ponce et al. (2019), colors highlight clustering into eight main cell types. Arrows indicate the direction of cell transitions which was estimated using RNA velocity. Red dots represent the possible neighborhood a cell can explore when modeled using a Markov chain. **B**. Propagator (Π) of the Markov chain as a function of the direction of differentiation (𝒱) and the stochasticity of gene expression (𝒟). Π_*ij*_ represents the probability of transitioning to the state *j* when being at the state *i*. **C-C’**. Stochastic differentiation trajectories starting in a Sox9 bipotent cell and ending in α (**C**), and β (**C’**) fates. **D-D’**. Fate probabilities calculated as the absorption probabilities for the α (**D**), and β (**D’**) sinks. **E-E’**. UMAP plots with normalized gene expression of known lineage- specific markers; Arx for α (**E**), and Nkx6-1 for β (**E’**) cells. **F-F’**. Average gene expression of known lineage-specific markers over α and β stochastic differentiation trajectories. **G-G’**. UMAP plot with motif activity profile of known lineage-specific regulators; Arx motif for α (**G**), and Nkx6-1 motif for β (**G’**) cells. **H-H’**. Average motif activity of known lineage-specific regulators over α and β stochastic differentiation trajectories. **I-I’**. Dynamic Pearson correlation between mRNA expression and motif activity over α and β trajectories; Arx and its motif (**I**), and Nkx6-1 and its motif (**I’**). **J-L**. Heatmaps showing the average motif activity over stochastic trajectories and dynamic Pearson correlation. Not all TF names are shown (see supplementary table 1 for full list). Right: activity distribution on the UMAP embedding of selected examples and respective z-score. **J**. Factors predicted as drivers of both β and α fates. **K**. Factors predicted as β-specific sorted according to which was active first in the β trajectories. **L**. Factors predicted as α-specific sorted according to which was active first in the α trajectories.

To retrieve the dynamic profiles towards the final endocrine fates, we first embedded the data in a low-dimensional manifold using UMAP with ten dimensions, and we built a nearest neighborhood graph in the reduced space. Next, we leveraged the robust RNA velocity profile to direct the edges of the Markov chain, Fig. 2B; and estimated transition probabilities using a velocity-driven kernel (see Methods). We used the transition matrix with a Monte Carlo sampling algorithm to simulate stochastic gene expression profiles along the differentiation trajectories, which allowed us to plot gene expression trends. The time-evolution simulation showed that, after two thousand iterations, the simulations mainly ended in three of the final endocrine fates α, β, and ϵ; with higher frequencies for α (12.4%) and β (77%), from now on we focused the analysis on these two lineages. In Fig 2C-C’, we presented an example of the simulated trajectories for the final fates α and β. These fates clearly represent sinks during the endocrine cell-subtype specification. We computed fate probabilities utilizing the information of stochastic simulations. We asked how often a simulated random walk that visited a given cell ended up in any terminal index sinks. We summarized this information in a sink-probability distribution (Fig 2D-D’). To verify the performance of our dynamic inference approach using different kernels, we also computed the transition probabilities with the gradient of potential energy as the drift. In this case we assumed that cells feel both repulsive and attractive forces with decreasing strengths as a function of their distances to progenitors and differentiated cells, respectively. As expected, we got similar sink-probabilities distribution (Supplementary figure 1). Of note, the inferred Fate probabilities follow the same trend of what has been reported for the same dataset using a deterministic approach ^26^, validating the subjacent dynamics of the stochastic simulations. Moreover, this analysis accurately identifies the β-fate as the most likely terminal fate for endocrine progenitors at E15.5, consistent with the previous biological knowledge ^17,27^.

Next, to evaluate the dynamic profile of transcriptional regulators, we estimated TF activities from the behavior of the predicted target genes. Briefly, there will be an increased activity for a TF when its targets show, on average, an increase in expression that cannot be explained by the presence of sites for other TFs in their promoters ^13^. To objectively assess the inferred transcriptional dynamics, we looked at known lineage-specific regulators’ expression and activity profiles. The TF Arx is essential for proper a cell formation ^28^, and unsurprisingly, it had high expression in α cells (Fig. 2E). As expected, the expression profile of Arx over the α trajectories increased while over β the trajectories was flat (Fig. 2F). Similarly, the TF Nkx6-1 acts downstream Neurog3 ^16^ and has been described as necessary for normal βcell development ^29^. Indeed, it was highly expressed in β cells (Fig. 2E’), and its expression profile increased only over β trajectories (Fig. 2F’). Strikingly, we found high Arx and Nkx6-1 activity in both α and β cells (Fig 2G-G’). We reasoned that this is the typical behavior of a TF that acts as an activator of one fate and repressor of the other. Indeed, a high activity means that the targets of the TF present a high expression on average, and the binding of other TFs in the promoters cannot explain this. Taken together, TFs that show correlation between its activity and its own mRNA expression are predicted to be activators. On the contrary, when the correlation is negative, a repression role is expected. We extracted the dynamical profile of Arx (Fig. 2H) and Nkx6-1 (Fig. 2H’) activities, and then performed dynamical correlation over α and β trajectories between the expression and the activity. We found that Arx is an activator of a α cell identity (positive correlation without any time-lag) while it is a repressor of the β lineage (negative correlation), Fig. 2I. This cell-dependent role has been well documented ^28,30^. On the other hand, Nkx6-1 behaves as an activator during cell β differentiation, whereas it has a repressor role in α cells, Fig. 2I’. This antagonistic behavior is supported by the findings of Schaffer et al. (2013) ^31^.

The differential motif activity analysis identified 86 TFs (Supplementary table 1), from which 22 were predicted to be specific for both α and β fates (Fig. 2J), 38 were β-specific (Fig. 2K), and 25 were α- specific (Fig. 2L). Interestingly, although Neurod1 was identified for both fates, the profile over the differentiation trajectories was higher in the β cells; this can also be observed in the distribution of activities in the UMAP plot (Fig. 2J). This result is consistent with previous publications about the role of Neurod1 in murine α- and β-cell specification, where the authors found a cell-type dependent role of Neurod1 in combination with Nkx2-2. They showed that Nkx2-2 represses Neurod1 in Pdx1+ and Neurog3+ progenitors allowing α-cell specification, while the activation of Neurod1 by Nkx2-2 permits β-cell formation ^32^. Although Nkx2-2 was not identified as differentially active, its activity profile is highly similar to the Neurod1 one (Supplementary figure 2). Bcl11a was the factor with the highest z-score (4.1192) (Fig. 2J), indicating the significance of this motif to explain the variance of the linear model of gene regulation. Remarkably, Bcl11a has an active role as a potent suppressor of insulin secretion in adult islets ^33^; however, despite its upregulation during the second wave of α-cell differentiation ^34^, its role during islet cell subtype specification remains to be studied. Some of the identified factors for both α and β fates have not been yet reported to have a function during pancreatic endocrine differentiation; thus, they are potential *novel* regulators. For instance, we found Cxxc1 as one of the differentially active factors early on during differentiation (Fig. 2J); it has already been pointed out as a critical factor during other differentiation processes such as in thymocyte development ^35^. Also, previous studies from our group reported it to be a direct target of Neurog3 ^36^.

Regarding the identified β-specific factors, our differential motif activity analysis identified several TFs known for playing a role in β-cell subtype specification or identity, including: Nkx6-1 ^29^, Glis3 ^37^, Mlxipl ^38,39^, and Pdx1 ^40^ (Fig. 2K). Noteworthy, Cebpg was the first factor in reporting an increasing activity profile over β-trajectories. It has been reported as a *novel* regulator of insulin secretion and transcription ^41^, whether it also has a function during β-cell differentiation remains to be unraveled. Similarly, among the factors designated as α-specific (Fig. 2L), we found known ones such as Arx ^28^ and Prox1 ^42^. Prox1 has a function during exocrine pancreas development ^43^, and it needs to be downregulated in β cells for their expansion and maturation ^42^. Hence, our predictions might suggest a role for Prox1 in α cell differentiation where this gene, in contrast to β cells, is highly expressed. We also identified core clock factors as Arntl as α -specific. Previous studies have shown that the distinct characteristics of α -cell and β-cell clocks harbor different circadian properties resulting in differential gene expression and functional regulation ^44^. Notably, one of the clock transcripts previously identified with an advanced phase in α -cells was Arntl ^44^. Intriguingly, Atf4 was the first factor to become active during α trajectories. It affects the pancreas morphology due to abnormal development of the acinar tissue ^45^ and has a paracrine effect on β cells ^46^. However, its effect on islet-cell subtype specification has not been studied. Thus, our predictions might indicate a cell-autonomous role of Atf4 in the α lineage. Taken together, FateCompass predicted well-known TFs, which serve as a positive control of the method’s performance. Beyond that, we identified *novel* potential β- and α - fate regulators with clearly distinct dynamic behaviors. This information can be harnessed further to characterize the regulatory interactions behind endocrine cell formation.

### FateCompass identifies transcriptional dynamic profiles beyond RNA velocity

To test whether the FateCompass workflow retrieves differentiation trajectories and identifies lineage-specific TFs in more complex experimental designs, with several harvesting points, we considered a scRNAseq experiment from an *in-vitro* differentiation of human stem cells towards pancreatic β –like cells ^18^. In this study, the authors used the SC-β-cell protocol to mimic β cell development ^47^. Shortly, human pluripotent stem cells growing in 3D clusters were differentiated into six stages using specific inducing factors to produce “stem-cell-derived islets” (SC-islets) that contained SC-β-cells (Supplementary Figure 3C). We applied FateCompass to a dataset of 25299 cells profiled using In-Drops sequencing across eight timepoints throughout stage five. Notably, at the beginning of stage five, there were NKX6-1+ pancreatic progenitors as well as the first SC-α cells; and by the end of it, there were three classes of endocrine cells: SC-β-cells expressing INS, NKX6-1, ISL1, PDX1, and other β-cell markers; SC-α -cells expressing GCG, ARX, IRX2 and also INS; and SC-EC-cells expressing CHGA, TPH1, LMX1A and SLC18A1 that resembled intestinal enterochromaffin (EC) cells (Fig. 3A). Noteworthy, enterochromaffin cells are not present, *in-vivo*, in mouse or human islets and are thus considered an *in-vitro*-specific fate bifurcation ^48,49^. We were interested in seeing how well FateCompass retrieved lineage-specific TFs in this challenging setting, where there is a directed differentiation (towards SC-islets) with an undesired by-product (SC-EC-cells).

**Figure 3.**
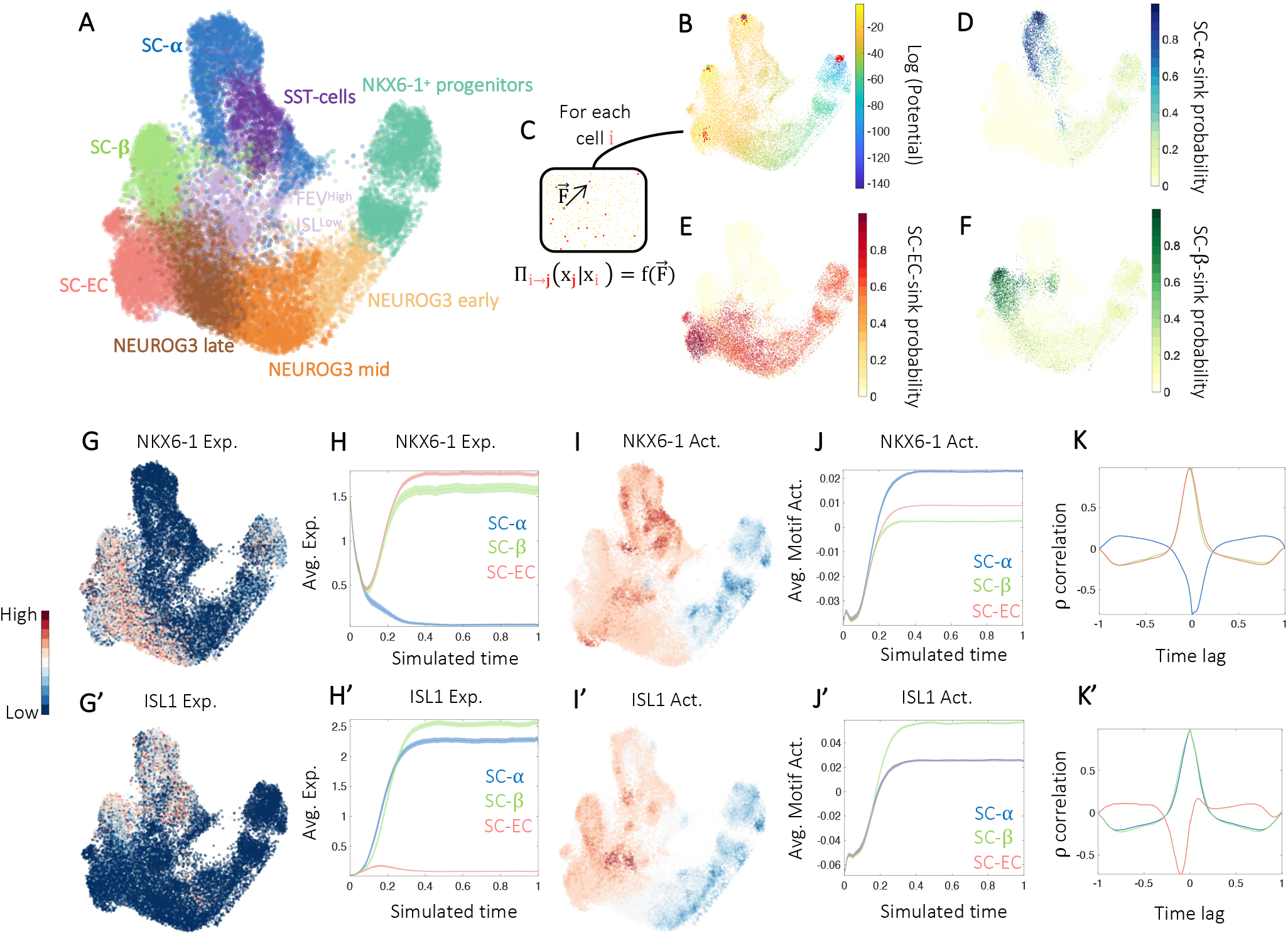
In-vitro β-cell differentiation landscape. **A**. UMAP plot of 25299 cells profiled during a 7-day time-course at stage 5 of differentiation towards β-like cells from Veres et al. (2019), colors highlight clustering into nine main cell types. **B**. UMAP plot colored according to the potential energy, the gradient goes from NKX6-1+ pancreatic progenitors (source) to the mature hormone- producing cell types (sinks). Red dots represent the possible neighborhood a cell can explore when modeled using a Markov chain. **C**. Propagator (Π) of the Markov chain as a function of the potential energy 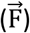 ∏_*ij*_ represents the probability of transitioning to the state *j* when being at the state *i*. **D-F**. Fate probabilities calculated as the absorption probabilities for the SC-α (**D**) SC-EC (**E**), and SC-β (**F**) sinks. **G-G’**. UMAP plots with normalized gene expression of known lineage- specific markers; NKX6-1 for SC-β and SC-EC (**G**) and ISL1 for SC-α and SC-β (**G’**) cells. **H-H’**. Average gene expression of known lineage-specific markers over SC-α, SC-β, and SC-EC stochastic differentiation trajectories. **I-I’**. UMAP plot with motif activity profile of known lineage-specific regulators; NKX6-1 motif for SC-β and SC-EC (**I**) and ISL1 motif for SC-α and SC-β (**I’**) cells. **J-J’**. Average motif activity of known lineage-specific regulators over SC-α, SC-β, and SC-EC stochastic differentiation trajectories. **K-K’**. Dynamic Pearson correlation between mRNA expression and motif activity over SC-α, SC-β, and SC-EC stochastic trajectories; NKX6-1 and its motif (**K**), and ISL1and its motif (**K’**).

To infer the differentiation trajectories towards the final endocrine cell types, we first computed RNA velocities using scVelo ^9^ and visualized them using 2-dimensional UMAP embedding (Supplementary Figure 3A). Notably, the projected velocities did not have a conclusive pattern towards the final fates, probably due to the high proportion of unspliced transcripts (30%, Supplementary Figure 3B). The above, together with the inherent limitation coming from the batch effect introduced by harvesting at different time points (Supplementary Figure 3C-D) ^10^, made the use of RNA velocity as a drift for the transition probabilities a liability. Therefore, we estimated FateCompass transition probabilities using the potential energy gradient from NKX6-1^+^ Progenitors to each terminal fate (SC-α, SC-β, and SC-EC), Fig. 3B-C (see Methods). To summarize the information of the stochastic simulations, we plotted the sink-probability distribution for each final cell type. Notably, the likelihood of having SC-α as final fate strongly decreases for NEUROG3-mid and NEUROG3-late progenitors suggesting that this cell type comes mainly from early endocrine precursors (Fig. 3D); this is consistent with previous reports ^50^. Oppositely, SC-β and SC-EC are the prevalent endpoints for trajectories passing through late NEUROG3 progenitors (Fig. 3E-F). These observations are consistent with previous studies ^18,51^, validating our drift-dependent Markov chain approach to infer differentiation dynamics.

To further validate FateCompass predictions, we checked the dynamic profile of known regulators. NKX6-1 is pivotal at different differentiation stages to giving rise to β-like cells ^50^. Congruently, Veres et al. (2019) reported high expression of NKX6-1 in early endocrine precursors, Neurog3-late progenitors, SC-β, and SC-EC cells (Fig 3G). In agreement, the dynamic profile of NKX6-1 expression (Fig. 3H) started at a high value that corresponds to the trajectories passing through early progenitors expressing NKX6-1+, PTF1A+, and PDX1^high^. Then, a decreasing profile is followed by an expected burst on SC-β and SC-EC cells, corresponding to the influence in endocrine cell-subtype specification ^18^. Similar to the *in-vivo* situation, we found high NKX6-1 activity in SC-β- and SC-α -; and in this context, also in SC-EC-cells (Fig. 3I-J). After checking the dynamical correlation of mRNA expression and motif activity (Fig. 3K), we consistently predicted NKX6-1 as an activator of the SC-β identity. Also, for the first time, we provide evidence of the possible role of NKX6-1 protein as an activator (positive correlation without time-lag with the mRNA expression in Fig. 3K) of the EC fate specification during pancreatic endocrine differentiation *in-vitro*. In contrast, the high NKX6-1 activity in SC-α cells that negatively correlated with the mRNA expression of the NKX6-1 transcript points to a possible repressor role in the SC-α-cells, suggesting a similar function to the reported by Schaffer et al. (2013) in the mouse ^31^. On the other hand, ISL1 is a well-known marker of β-cells ^52^, and it functions as a regulator of ARX during α-cell development ^53^. Indeed, Veres et al. (2019) reported it as differentially expressed in the SC-β branch, Fig. 3G’; and the dynamic expression profile showed a clear increasing pattern over SC-α and SC-β differentiation trajectories (Fig. 3H’). Moreover, the activity profile of the ISL1 motif was higher in the expected populations, SC-α- and SC-β-cells (Fig. 3I’-J’). Also, the dynamic correlation between expression and activity (Fig. 3K’) confirmed the expected activator role over SC-α and SC-β trajectories. Altogether, these results validate both the potential energy from progenitors to final fates as a useful tool to reveal differentiation trajectories and the linear model of gene regulation as an efficient way to estimate TF activities in challenging systems.

### Differential motif activity analysis predicts driving factors during *in-vitro* -cell differentiation protocols

To check FateCompass performance in identifying lineage-specific regulators on a challenging differentiating system, whose endpoints include an off-target population, we applied the differential motif activity analysis to the *in-vitro* differentiation towards pancreatic β-like cells dataset. We identified 126 differentially active TFs (Supplementary Table 2), 14 for the three endocrine lineages (Fig. 4A), 14 for both SC-β and SC-α (Fig. 4B), 15 for SC-β and SC-EC (Fig. 4C), 10 for SC-α and SC-EC, 20 were SC-β-specific (Fig. 4D), 25 were SC-EC-specific (Fig. 4E), and 28 were SC-α -specific (Fig. 4F). Interestingly, the TF CDX2 was differentially active for SC-β, SC-EC, and SC-α; this finding was puzzling since CDX2 is well-known for its role in intestinal specification of the gut endoderm during development ^54^. During mammalian development, different organs such as the stomach, pancreas, liver, and intestine derive from the gut endoderm. Different regulatory interactions control gut endoderm regionalization promoting organ specification. It has been reported that the TFs PDX1 and SOX9 have a positive cross-regulatory loop that promotes the expression of pancreas-specific factors while repressing CDX2 ^55^. Thus, our predictions suggest that the early endodermal progenitors might still be plastic and have the potential to activate other fates that will be repressed upon islet-cell development; this observation agrees with the reported by Ramond et al. (2018) ^56^. Of note, CDX2 activity is high at the beginning of the three endocrine fates trajectories, then, it decreased in SC-β and SC-α cells while remaining high in SC-EC, which resemble intestinal enterochromaffin cells (Fig. 4A). For its part, MAFB was also identified for the three endocrine fates. However, its activity was higher through SC-β and SC-α trajectories (Fig. 4A); this predicted behavior agrees with the known role of MAFB during islet α and β cell development ^57^.

**Figure 4.**
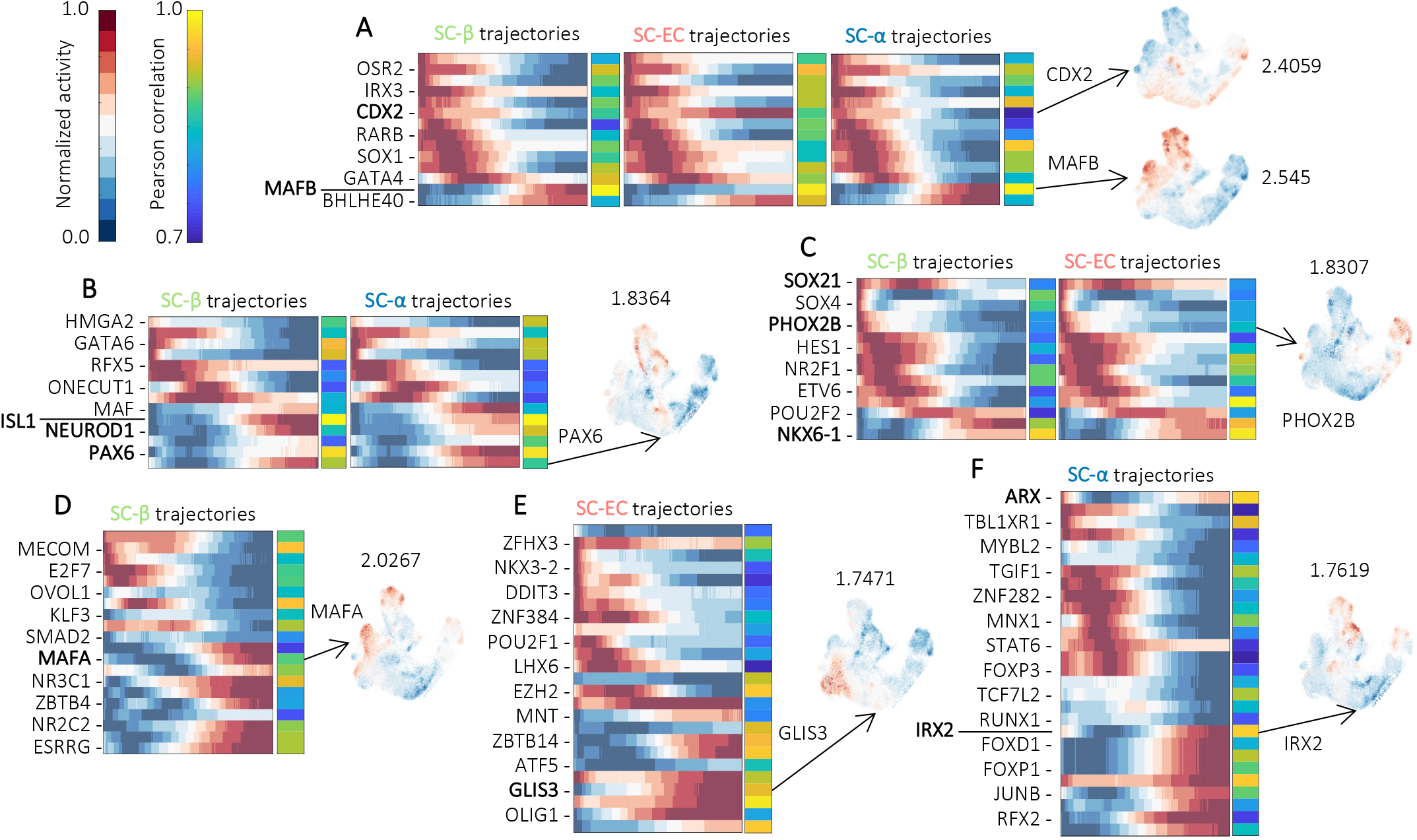
Differential motif activity analysis during in-vitro β-cell differentiation. Heatmaps showing the average motif activity over stochastic trajectories and dynamic Pearson correlation. Not all TF names are shown (see supplementary table 2 for full list). Right: activity distribution on the UMAP embedding of selected examples and respective z-score. **A**. Factors predicted as drivers of the three lineages: SC-β, SC-EC, and SC-α. **B**. Factors predicted as drivers of both SC-β and SC-α fates. **C**. Factors predicted as drivers of both SC-β and SC-EC fates. **D**. Factors predicted as SC-β- specific sorted according to which was active first in the SC-β trajectories. **E**. Factors predicted as SC-EC-specific sorted according to which was active first in the SC-EC trajectories. **F**. Factors predicted as SC-α-specific sorted according to which was active first in the SC-α trajectories.

FateCompass predicted some factors to be specific for two lineages simultaneously. As expected, ISL1 and NEUROD1 were classified as SC-β- and SC-α-specific (Fig. 4B) ^32,52,53^. Contrary to what we observed in the mouse *in-vivo* situation, where NEUROD1 and NKX2-2 had a similar activity profile, in this case, during human *in-vitro* β-cell differentiation, the activity of NKX2-2 was restricted to NEUROG3 endocrine progenitors and SC-EC cells (Supplementary figure 4). This observation points to differences in the regulatory programs of endocrine cell differentiation in mice vs. humans. PAX6 was also identified as SC-β- and SC-α-specific (Fig. 4B). Previous chromatin analysis and shRNA-mediated gene suppression experiments showed that PAX6 has a key role in the identity and function of β-cells by activating specific markers and repressing alternative islet genes. Interestingly, using RNAseq and luciferase assay, the authors found that PAX6 represses NKX2-2 ^58^. We observed mutually exclusive behavior for PAX6 and NKX2-2 activities (Supplementary figure 4), which supports that PAX6 and NKX2-2 might have antagonistic roles. Regarding the factors classified as SC-β- and SC-EC-specific (Fig. 4C), we found NKX6-1 among the predictions, supporting our previous observation of the possible activator role of NKX6-1 for both the pancreatic β-like cells and the off-target intestinal-like population of EC cells. Along the same line, a recent study found an enrichment of the NKX6-1 motif on EC-like cells using single-cell ATAC seq ^59^, which further endorse our prediction. Notably, we identified PHOX2B, which belongs to the same motif family as LMX1A (Supplementary figure 5); LMX1A has a known role as a regulator of the EC fate in the adult small intestine downstream NKX2-2 ^60^. Thus, our findings suggest that promoters with the binding site for PHOX2B/LMX1A are, on average, highly expressed on the SC-β- and SC-EC-trajectories. Whether the differentiation of SC-EC cells has similar regulatory mechanisms to those in the murine small intestine remains largely elusive.

To learn more about SC-β, SC-α, and SC-EC development, we exclusively focused on FateCompass predictions for each particular lineage. MAFA was among the SC-β-specific factors (Fig. 4D), indicating that in the human *in-vitro* context, there is a similar switch from MAFB to MAFA, in developing towards fully functional β-cells, to the one reported in rodents ^61^. GLIS3, which we previously found as a direct target of NEUROG3 in pancreatic endocrine progenitor cells ^36^, was classified as SC-EC (Fig. 4E). Notably, it has been reported both as a β-cell marker in the pancreas ^37^ and as an EC marker in the adult small intestine ^62^. Thus, our data might imply a tissue- and cell-dependent role for GLIS3. ARX is the first TF to become highly active during SC-α trajectories, corroborating its function during glucagon-producing cells development ^28^. IRX2 was highly active later on during SC-α specification (Fig. 4F); importantly, Gage et al. (2015) found it downregulated in hPSC-derived human islet cells lacking ARX ^63^, and Schreiber et al. (2021) found it as a direct target of NEUROG3 ^36^. Hence, our predictions and previous evidence position IRX2 as a potential α-specific novel TF acting downstream NEUROG3. Taken together, FateCompass systematically predicts known and novel potential regulators during a complex differentiation system that, besides the expected population, included an off-target cell type, highlighting the possible use of the pipeline in the improvement of differentiation protocols.

### Comparison between *in-vivo* and *in-vitro* β-cell-specific drivers

The design of stepwise directed differentiation protocols to produce islet-like cells has relied heavily on mouse pancreas developmental biology knowledge. We leveraged the predictions of FateCompass to identify common and new transcriptional interactions to steer hypothesis generation aiming to unravel what leads to the cellular heterogeneity of β cell differentiation protocols. First, we reasoned that by comparing the differentially active factors involved in β-cell specification, we could get insights into similar and different regulatory programs in mouse *in-vivo* and human *in-vitro*. We found an interesting pattern of conservation and divergence with only around 16% of TFs at the intersection (Fig. 5A). We reasoned that having few overlapping factors could be due to significant differences at the expression level that translate in different TFs driving such an expression pattern. We performed hierarchical clustering among the mouse *in-vivo* and human *in-vitro* populations to test this. We found that mouse β-cells are more similar to the rest of murine hormone-producing cells (α, ϵ, and δ), with a Pearson correlation higher than 0.7 (Supplementary figure 6). Although human-derived SC-β cells clustered with most of the endocrine-committed murine cells, their relationship with mouse β-cells was not that high, Pearson correlation 0.6 (Supplementary figure 6). The above points to significant differences in the expression levels that drive the inference of mainly divergent TFs.

**Figure 5.**
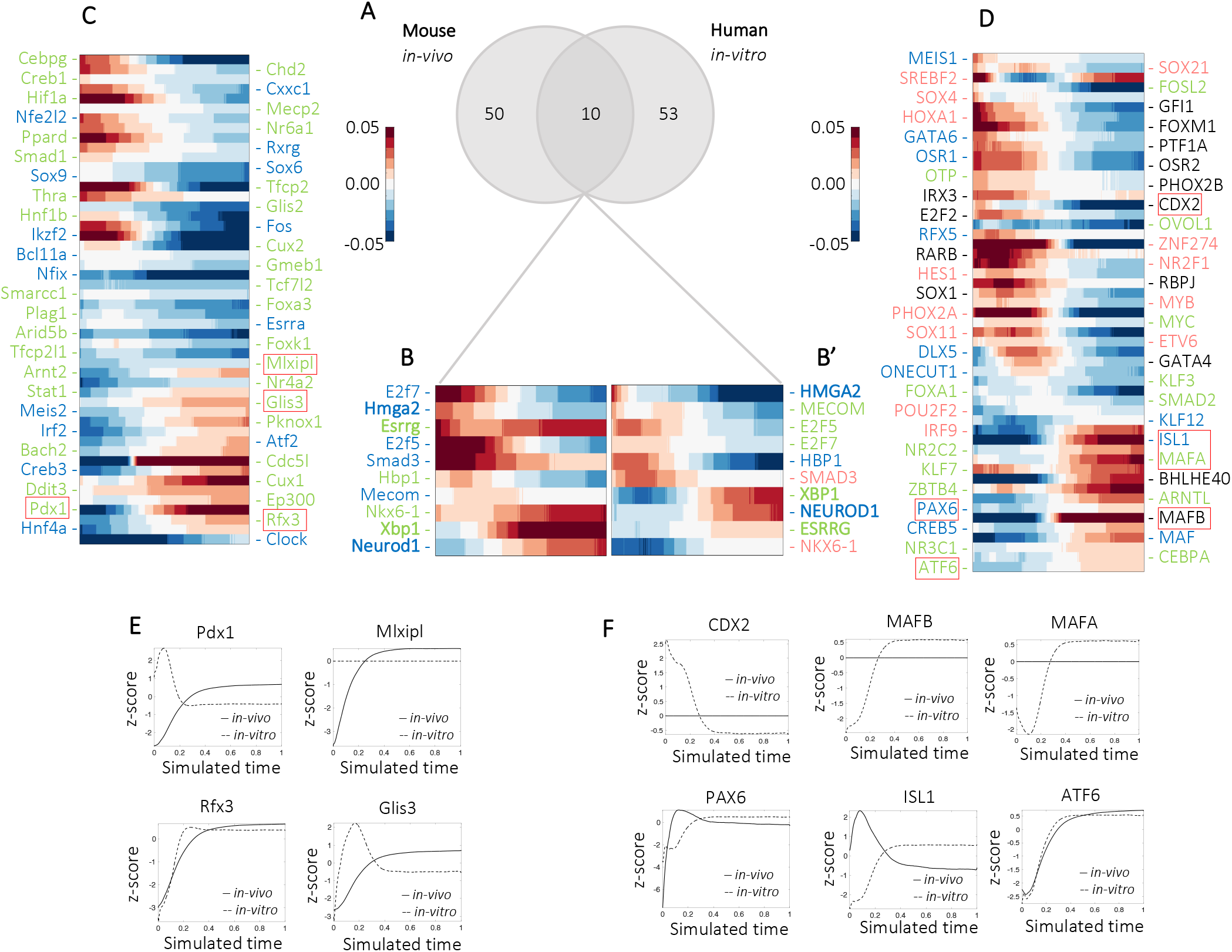
Cross-species comparison of the β-specific factors. **A**. Venn-diagram showing the number and overlap of β-specific TFs predicted by FateCompass. **B-B’**. Heatmaps showing the average motif activity over β trajectories of the 10 overlapping factors in the mouse *in-vivo* system (**B**), and in the human *in-vitro* system (**B’**). **C**. Heatmaps showing the average motif activity over β trajectories of the 50 mouse *in-vivo* specific factors, TFs are sorted according to which was active first. **D**. Heatmaps showing the average motif activity over β trajectories of the 53 human *in-vitro* specific factors, TFs are sorted according to which was active first. **E-F**. z-score of the average motif activity profile over β trajectories for some known factors specifically predicted for the mouse *in-vivo* dataset (**E**) and for the human *in-vitro* dataset (**F**). <u>Color code for the name of the TFs in the heatmaps</u>: green for the β-specific, blue for the β- and*α*-specific, pink for the β- and EC-specific, and black for the β-, *α*- and EC- specific.

Next, we focused on the common TFs for the mouse *in-vivo* and the human *in-vitro* to see whether there were relative-time-specific profiles. To that end, we sorted the TFs according to which was active first for each system independently (Fig. 5 B-B’). Hmga2 was classified, in both cases, as β- and α- specific, and it was active at early stages. Conversely, Essrg had a different dynamic profile despite being β-specific in both systems. While it was constantly highly active in the mouse *in-vivo* and appeared third on the dynamic ranking, it had a slowly increasing profile in the human *in-vitro* and showed next to last. This result was striking because the estrogen-related receptor γ (Esrrg) is a hallmark of adult and not developing β-cells, with a known function for metabolic maturation ^64^. Therefore, we did not expect a high activity on the mouse system. Then, our prediction might indicate that Esrrg has a stage-dependent role that remains to be explored in the embryo. Notably, Neurod1 was β- and α-specific, appearing at later stages in both contexts, indicating that Neurod1 is playing a similar role in both systems. Similarly, Nkx6-1 was a common factor that became active progressively during the β-differentiation trajectories. Notwithstanding, Nkx6-1 was also involved in the EC cell subtype specification in the human *in-vitro* dataset, which raises flags about its organism-dependent role and the different programs it is activating.

To unravel possible time-specific regulatory interactions, we plotted the dynamic profiles of the species-specific TFs (Fig. 5C-D). About the mouse-specific factors (Fig. 5C), the dynamic profiles showed that Sox9 was active early on, confirming its role in inducing Neurog3 in the progenitor cord ^65^. Additionally, we found Meis2 with an increasing profile towards the end of the β-trajectories; previous studies reported it to be enriched in the second wave of murine fetal α cells ^66^, but its specific role in β-cell differentiation remains known. We observed two dynamic waves regarding the human-specific factors (Fig. 5D). The first includes factors active early on during the SC-βtrajectories with fetal-like functions, such as SOX4, PTF1A, and CDX2; the second with factors active later on during β-cell differentiation resembling maturation and maintenance roles, such as MAFA and PAX6. The above suggests that the human *in-vitro* regulatory programs differentiate β-like cells activating adult-like factors to produce functional insulin-responsive cells. Interestingly, FateCompass identified GATA4 as human-specific; this factor represents a well-known human-mouse difference. Indeed, its expression is delayed during human development, appearing simultaneously as PDX1 ^67^. Next, to learn more about differences in some known factors identified for one system and not the other, we plotted the z-score of their average activity profile over β-trajectories for the mouse *in-vivo* vs. the human *in-vitro* (Fig. 5E-F). Naturally, TFs identified only in the mouse system such as PDX1, MLXIPL, RFX3, and GLIS3 presented a constantly increasing profile in the *in-vivo* dataset. In contrast, in the *in-vitro* dataset, they either had a burst and then decreased or were not active (Fig. 5E). Similarly, except for CDX2, known β-cell markers predicted only in the human case (MAFB, PAX6, ISL1, MAFA, and ATF6) presented an increasing profile in the *in-vitro* dataset. At the same time, in the *in-vivo* system, no clear pattern was identified for these TFs (Fig. 5F).

Altogether, comparing the mouse *in-vivo* vs. human *in-vitro* β-cell differentiation showed that significant differences in gene expression profiles lead to inferring different lineage-specific TFs. Also, FateCompass recovered robust well-known TFs at the intersection of both systems. Beyond the similarities, we found that factors involved in early development prevailed in the mouse system. In contrast, a second dynamic wave that activates factors involved in maturation and maintenance is also present in the human case.

### FateCompass guides hypothesis generation to understand SC-EC cell-fate determination during *in-vitro* β-cell differentiation protocols

To further investigate the possible role of NKX6-1 in differentiating the undesired SC-EC cells during *in-vitro* β-cell differentiation, we looked at the targets of the NKX6-1 motif in the mouse and the human (Fig. 6A). Remarkably, CDX2, a small intestinal epithelial marker ^68^, and NKX2-2, a well-documented factor for having a role in developing endocrine cells in the small intestine and the pancreas, are human-specific targets. In the mouse developing pancreas, the expression of Nkx2-2 starts at E12.5, and it promotes α and β fates while repressing ghrelin-producing cells ^69^; at later stages (E15.5 and E18.5), the knockout of Nkx2-2 downstream Neurog3 results in defective β-cell differentiation ^70^. On the other hand, in the developing small intestine, Nkx2-2 expression starts at E15.5, and serotonin, the main secretory product of EC cells, is reduced in the absence of Nkx2-2 ^71^. Similarly, in the adult small intestine, the role of Nkx2-2 downstream Neurog3 regulating Lmx1a, a direct regulator of Tph1 (enzyme involved in the synthesis of serotonin), has been previously proven ^60^. Altogether, the mentioned studies about Nkx2-2 in the mouse suggest it has different roles: tissue- and stage-specific. Of note, in the mouse *in-vivo* dataset, the expression of Nkx2-2 was very sparse and mainly in progenitor cells, and activity of the Nkx2-2 motif was restricted to the progenitor and pre-endocrine cells (Supplementary Figure 2). In the human *in-vitro* dataset, NKX2-2 expression was high from Neurog3+ clusters to SC-β and SC-EC cells, and its motif activity was high from Neurog3+ cells towards SC-EC cells, not towards SC-β cells (Supplementary Figure 4). Then, we reasoned that NKX6-1 motif activity and NKX2-2 mRNA level should have a similar pattern if there is direct activation. Similarly, NKX2-2 motif activity should follow a similar trend to the LMX1A mRNA level. We found an increasing, almost parallel, profile for NKX6-1 activity and NKX2-2 expression (Fig. 6B). On the other hand, NKX2-2 activity peaked when the LMX1A expression level started to increase (Fig. 6C), which might indicate an early burst of activation.

**Figure 6.**
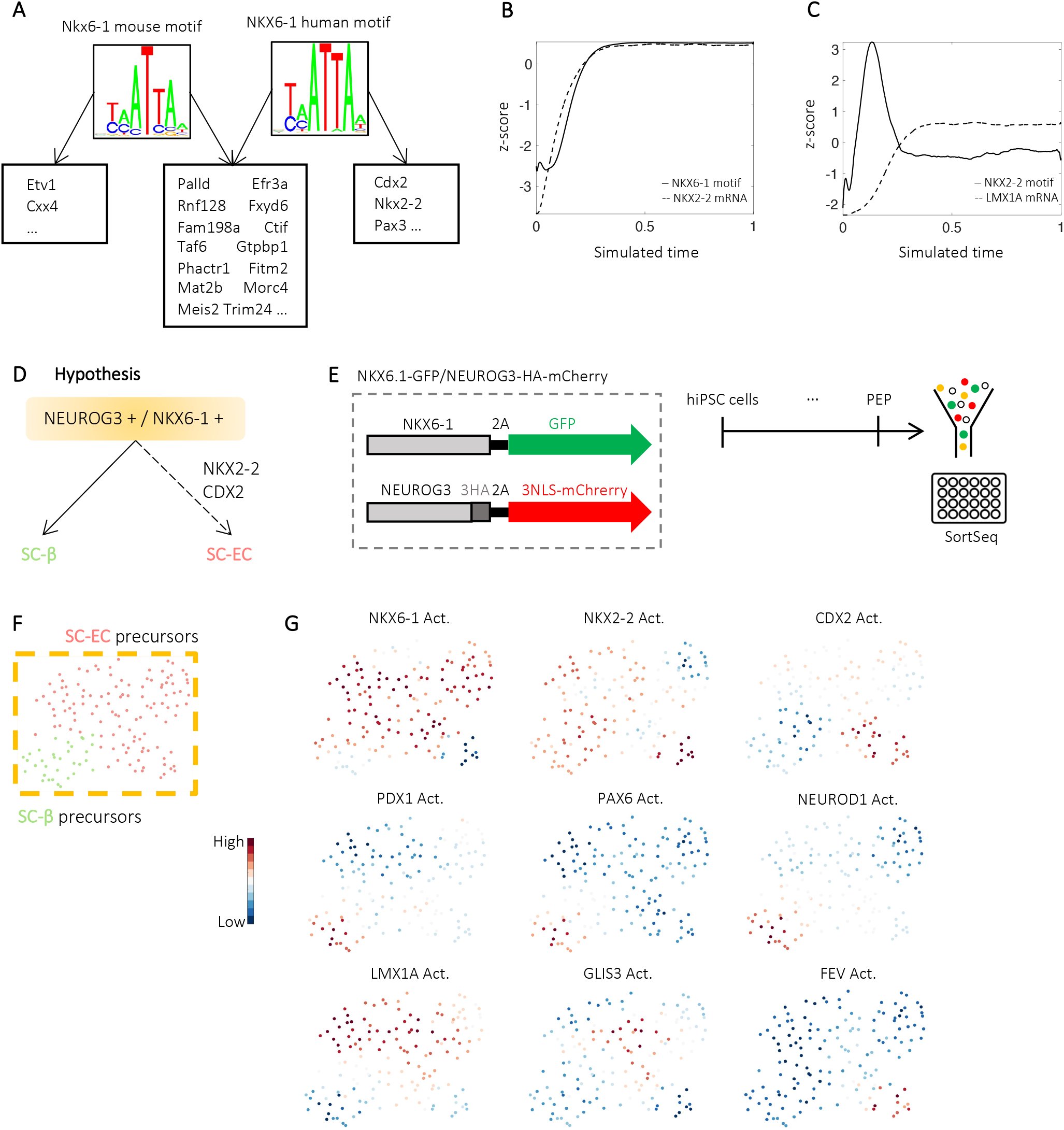
Role of NKX6-1 on the differentiation of the off-target population SC-EC. **A**. Nkx6-1 common and diverging targets for the mouse and human motif. **B**. z-score of NKX6-1 average motif activity and NKX2-2 average gene expression over SC-EC trajectories. **C**. z-score of NKX2-2 average motif activity and LMX1A average gene expression over SC-EC trajectories. **D**. Diagram of the hypothesis generated using FateCompass predictions. **E**. Scheme of the experimental design to test the generated hypothesis. NKX6.1-GFP/NEUROG3-HA-mCherry cell line to lineage trace the NEUROG3+ and NKX6-1+ population. Cell differentiation, sorting, and sequencing strategy. **F**. UMAP plot of 150 cells from the double-positive population. **G**. UMAP plot with motif activity profile of lineage-specific regulators.

To validate the hypothesis generated with FateCompass predictions on the mechanisms of SC-EC generation (Fig. 6D), we lineage-traced NEUROG3-NKX6-1positive cells at the pancreatic endocrine progenitor (PEP) stage and examined the heterogeneity of this population. To that end, we first generated an hiPSC line where NEUROG3 is fused to a cleavable mCherry reporter and NKX6-1 to a GFP reporter (NKX6.1-GFP/NEUROG3-HA-mCherry) (Fig. 6E). Next, we differentiated the NKX6.1-GFP/NEUROG3-HA-mCherry hiPS cells towards the islet lineage until the PEP stage at day 13, FACS sorted the differentiated cells and sequenced the different fluorescent populations using the single-cell RNA sequencing method SortSeq ^49^ (Fig. 6E and Supplementary Fig. 7A-B). The sequenced cells clustered according to the reporter for which they were sorted (Supplementary Fig. 7C). The negative cells expressed pluripotency markers such as SOX17; the GFP+ cells were more endodermal progenitors expressing SOX9 and CXCR4; and, the mCherry+ cells were endocrine committed cells expressing CHGA, GCG, INS, etc. We focused on the double-positive population, which clearly expressed markers of both SC-β cells and SC-EC cells (Supplementary Fig. 7D). Indeed, after clustering the double-positive population, we identified two groups SC-β precursors and SC-EC precursors (Fig 6F and Supplementary Fig. 7 E-F). This confirmed that SC-β and SC-EC cells differentiate from similar progenitor cells after pancreatic endocrine commitment downstream NEUROG3 and NKX6-1. Furthermore, we computed motif activities to see the potential regulators of each fate. We found NKX6-1 activity in both SC-β precursors and SC-EC precursors; NKX2-2 followed a similar pattern, whereas CDX2 activity was higher only in SC-EC precursors (Fig. 6G). SC-β-known drivers such as PDX1, PAX6, and NEUROD1 had high activity on the SC-β precursors. Likewise, the well-described enterochromaffin markers LMX1A and FEV were highly active in the SC-EC precursors; interestingly, GLIS3, a factor that FateCompass predicted to be a driver of the SC-EC fate, also had high activity on the SC-EC precursor cells (Fig. 6G).

Taken together, by exploring the behavior of common TFs and digging further into the transcriptional interactions of NKX6-1, we opened the question of whether the SC-EC differentiation is regulated by the TFs NKX2-2 and CDX2 acting downstream NKX6-1. We tested this hypothesis by lineage-tracing NEUROG3/NKX6-1 positive cells and found a population of precursors with clear SC-β and SC-EC characteristics. Interestingly, CDX2 activity was high exclusively on the SC-EC-precursors indicating that promoters with a binding site for CDX2 were, on average, highly expressed in this population. This type of hypothesis merits further investigation to identify key targets to improve β-cell differentiation protocols.

## Discussion

Here, we have introduced FateCompass, a workflow that robustly estimates lineage-specific TFs dynamically. FateCompass pipeline integrates a flexible framework to infer gene expression dynamic profiles with a linear model of gene regulation based on interactions between TFs and promoters to predict regulators implicated in fate choice during development in different contexts (*in-vivo* and *in-vitro*), across sequencing platforms (10X and InDrops) and across organisms (mouse and human). We designed an innovative differential motif activity analysis that considers the significance of the TF to explain the variability of the linear model of gene regulation, the change of the regulatory activity throughout the cell-fate decision process, and the dynamical correlation of the TF activity with the TF mRNA level. Applied to pancreatic islet cell subtype specification, we predicted time-and fate-specific known and novel TFs; the former serve as ground truth while the latter represents an advance in the current understanding of the transcriptional interactions underlying endocrine cell differentiation.

In the inference of differentiation trajectories, we assumed, like other studies, that the process of a cell changing states along a trajectory until it reaches a final fate can be understood as a particle diffusing on a volume ^7,72^; but, unlike them, we infused the direction of the differentiation as a drift to bias the transition probabilities. When RNA velocity profiles are robust, FateCompass uses them to direct the edges of the Markov chain. On this line, FateCompass predicted the same terminal fates and final fate probabilities for murine endocrine differentiation as similar existing methods ^26^. During development, when the starting cell and the final fates are clear, and the RNA velocity profiles are inconclusive, FateCompass infers differentiation trajectories beyond RNA velocity, biasing the transition probabilities using the gradient of potential energy from the starting cells towards the final states. We validated this approach using a dataset from an *in-vitro* differentiation towards β-cells, where we accurately recovered that β-like cells differentiate from NEUROG3-late progenitors while α - like cells start to differentiate from NEUROG3-early progenitors ^18,50,51^.

FateCompass uses TFs as the leading players in the gene regulation model; they are well-known for their direct role in gene-specific transcriptional regulation; hence they are commonly used as readouts of pathway activities ^73^. Other approaches attempting TF activity inference from transcriptomic data, both bulk and single-cell, do not consider the dynamic nature of the TF activities ^2,14,15,74^. Some studies have based their predictions merely on correlations between mRNA level of the TF and expressed genes ^15,74^. Other more advanced rely on known regulons and inferred TF activities using the correlation of the mRNA level of the TF and the group of genes that it can potentially regulate, based on the presence of binding sites on a given regulatory region ^2,14^. In contrast to the previously cited methods, ISMARA, initially developed for bulk RNA seq data, does not rely on correlations; it modeled the expression levels as a linear combination of TF binding site predictions and unknown TF activities ^13^. Here, we extended the use of ISMARA to single-cell transcriptomics. The original ISMARA model proposed a symmetric Gaussian prior to avoid overfitting; however, in that way, all the parameters are regularized equally, which might not be suitable on single-cell data, where different regions of the manifold represent, usually, different phenotypes associated with changing TF activities. FateCompass address the multicollinearity problem in linear regression using a novel regularization approach. We defined a data-diffusion-based regularization where we enforced the smoothness and stability of the inferred activities across cells. This approach has been widely used for imputation methods ^25^.

In the mouse *in-vivo* data set, FateCompass recovered well-documented regulatory interactions, such as the antagonistic role of Arx and Nkx6-1 ^30,31^ and the cell-type dependent interaction between Neurod1 and Nkx2-2 ^32^. We also identified putative driver factors with interesting known roles in β cell function ^41^ and a circadian pattern in α cells ^44^, whether they are also involved in endocrine cell differentiation remains to be tested. A recently published study aimed to identify lineage-specific drivers during pancreatic endocrine differentiation, where they focused on the differential gene expression of TFs, also identified some of the known regulators ^75^. In contrast, we steered on regulation principles by considering interactions between TFs and promoters, which provide a more accurate picture of gene-specific regulation. We anticipate applying our framework to guide experiment design to test the function of the lineage-specific factors. In the human *in-vitro* dataset, we identified TFs acting early on during the differentiation trajectories that confirmed the plasticity of the less mature cells in differentiation protocols ^56,68^. Moreover, we retrieved cell-type-specific drivers for the pancreatic endocrine cells and the off-target enterochromaffin population. Importantly, our differential motif activity analysis pinpointed, for the first time, NKX6-1 as a potential regulator of the SC-EC cells, which was further experimentally validated by generating single-cell transcriptomes of lineage-traced cells. Comparing the *in-vivo* and *in-vitro* predictions for the β cell trajectories, we found known TFs at the intersection such as NEUROD1 and NKX6-1 ^31,32^; also, we were able to recapitulate mouse- and human-specific differences ^67^. In summary, we foresee the use of FateCompass to generate hypotheses targeted to provide means to optimize differentiation protocols.

The fast evolution of high-throughput methods and generation of large-scale datasets impose the need for robust computational approaches not only to characterize genome-wide patterns but also to extract information and mechanistically model biological phenomena that, in the end, will provide predictions aimed at increasing the current state of the knowledge. As with any inference method, aspiring to reconstruct the exact interactions underlying a complex biological process, such as endocrine cell formation, is a futile task. In this study, we rely on computationally predicted regulatory sites, summarized in a binding site matrix; this represents a bias on the structure of the gene regulatory network. Moreover, we are only considering interactions between TFs and promoters, and it is well known that some essential regulatory interactions occur at distal regulatory sites ^76^. We have designed our pipeline such that the limitations mentioned above could be addressed by extending the binding site matrix. As a framework for identifying lineage-specific drivers, we forecast FateCompass to be used as a tool to explore scRNAseq data, guide hypothesis generation, and direct experiment design. Further experimental validation of the generated hypothesis will increase the current understanding of a given process and provide means to improve existing translational experiments aimed at cell therapy.

## Acknowledgements

We thank E. van Nimwegen and G. la Manno for stimulating discussions on pipeline development; L. McInnes for valuable insights on UMAP implications, interpretation, and limitations. We thank Christian Honoré (Novo Nordisk A/S) for providing the NKX6.1-GFP iPSC line ^77^ through the IMI/EU sponsored StemBANCC consortium. We would further like to thank the IGBMC Flow cytometry facility for cell sorting and the members of the Gradwohl lab and the Molina lab. Research in the Gradwohl and Molina lab is supported by a grant from ANR (ANR-21-CE14-0003-01). IdEx Unistra [ANR-10-IDEX- 0002]; SFRI-STRAT’US project [ANR 20-SFRI-0012]; EUR IMCBio [ANR17-EURE-0023]. Sara Jiménez is an IGBMC international PhD programme fellow supported by the LaxEx INRT.

## Author contributions

S.J., G.G. and N.M. conceptualized the project. S.J. and N.M. developed and implemented the method. G.G., V.S. and S.J. interpreted the relevance of the method for inferring lineage-specific pancreatic endocrine drivers. V.S. and R.M. performed hiPSC gene editing, differentiations and cell sorting. G.G. and N.M supervised the project. S.J. analyzed the data and wrote the manuscript with contribution of all coauthors. All authors read and approved the final manuscript.

## Data and Code availability

Single cell RNA sequencing data of this study have been deposited in the Gene Expression Omnibus (GEO) under accession code GSE202092. Scripts to reproduce our analysis are available at https://github.com/sarajimenez/fatecompass

## Supplemental information

Supplementary figures: Supplementary Figures 1-7.

Supplementary tables: Supplementary tables 1-2.

---

**Supplementary figure 1.**
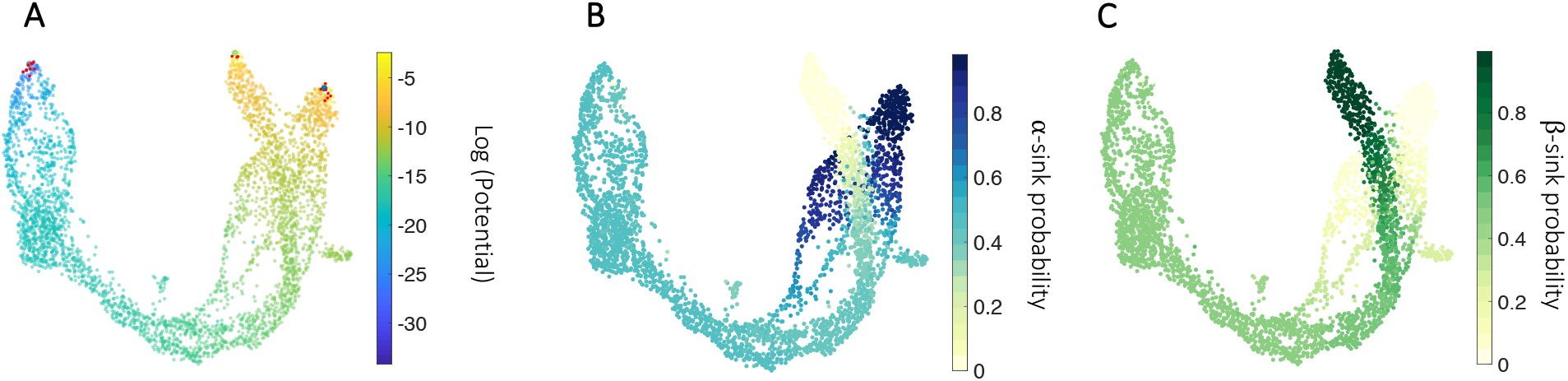
Inference of differentiation trajectories using potential energy in the mouse system. **A**. UMAP plot colored according to the potential energy, the gradient goes from Sox9+ bipotent progenitor cells (source) to the mature hormone-producing cell types (sinks). Red dots represent the possible neighborhood a cell can explore when modeled using a Markov chain. **B-C**. Fate probabilities calculated as the absorption probabilities for the α (**B**), and β (**C**) sinks.

**Supplementary figure 2.**
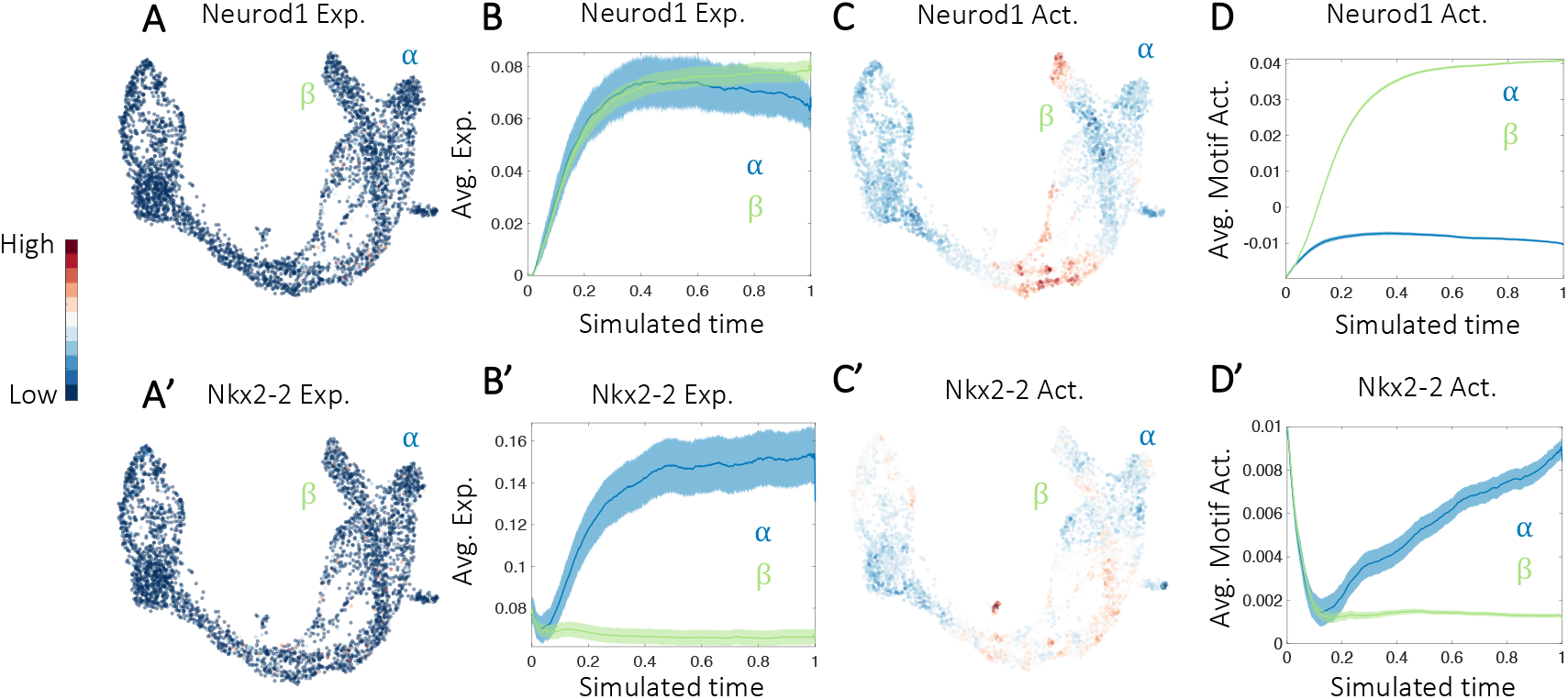
Expression and activity profile of known regulatory interaction between Neurod1 and Nkx2-2 in the mouse in-vivo data set. **A-A’**. UMAP plots with normalized gene expression of Neurod1 (**A**), and Nkx2-2 (**A’**). **B-B’**. Average gene expression over α and β stochastic differentiation trajectories for Neurod1 (**B**), and Nkx2-2 (**B’**). **C-C’**. UMAP plot with motif activity profile of Neurod1 motif (**C**), and Nkx2-2 motif (**C’**). **D-D’**. Average motif activity over α and β stochastic differentiation trajectories for Neurod1 (**D**), and Nkx2-2 (**D’**).

**Supplementary figure 3.**
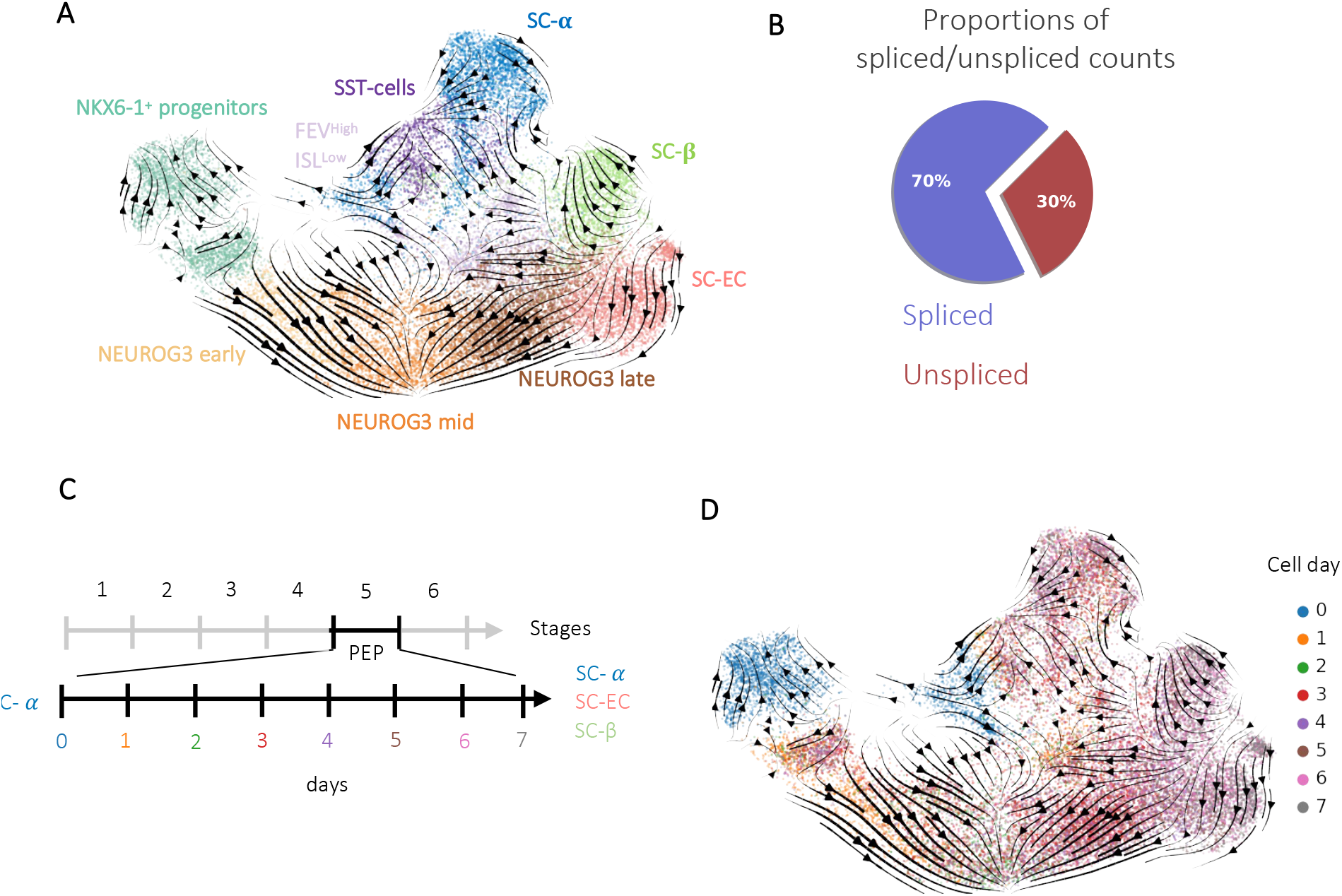
RNA velocity pitfalls in a slow-differentiating system. **A**. UMAP plot of 25299 cells profiled during a 7-day time-course at stage 5 of differentiation towards β-like cells from Veres et al. (2019), colors highlight clustering into nine main cell types, and arrows indicate the future state of the cells predicted with the dynamical model of RNA velocity, scVelo. **B**. Proportions of spliced/unspliced counts in the data set. **C**. Summary scheme of the differentiation stages towards β-like cells and zoom into stage 5, where, in the beginning, there are the Pancreatic Endocrine Progenitors (PEP), and at the end, there are hormone-producing cells. **D**. UMAP plot colored by the cell day in which they were sampled.

**Supplementary figure 4.**
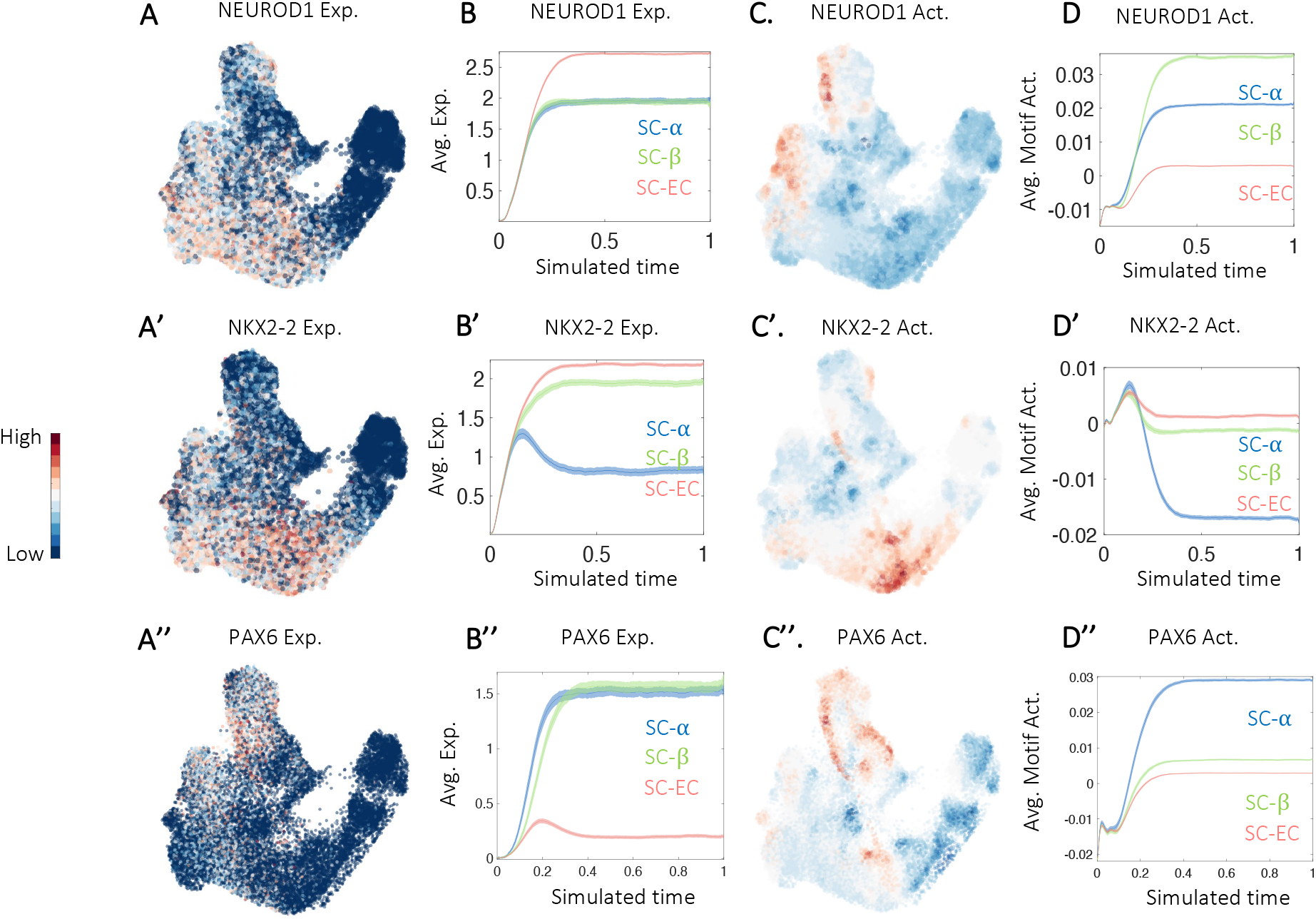
Expression and activity profile of known regulatory interaction between NEUROD1, NKX2-2, and PAX6 in the human in-vitro data set. **A-A’’**. UMAP plots with normalized gene expression of NEUROD1 (**A**), NKX2-2 (**A’**), and PAX6 (**A’’**). **B-B’’**. Average gene expression over SC-α, SC-β, and SC-EC stochastic differentiation trajectories for NEUROD1 (**B**), NKX2-2 (**B’**), and PAX6 (**B’’**). **C-C’’**. UMAP plot with motif activity profile of NEUROD1 motif (**C**), NKX2-2 motif (**C’**), and PAX6 motif (**C’’**). **D-D’’**. Average motif activity over SC-α, SC-β, and SC-EC stochastic differentiation trajectories for NEUROD1 motif (**D**), NKX2-2 motif (**D’**), and PAX6 motif (**D’’**).

**Supplementary figure 5.**
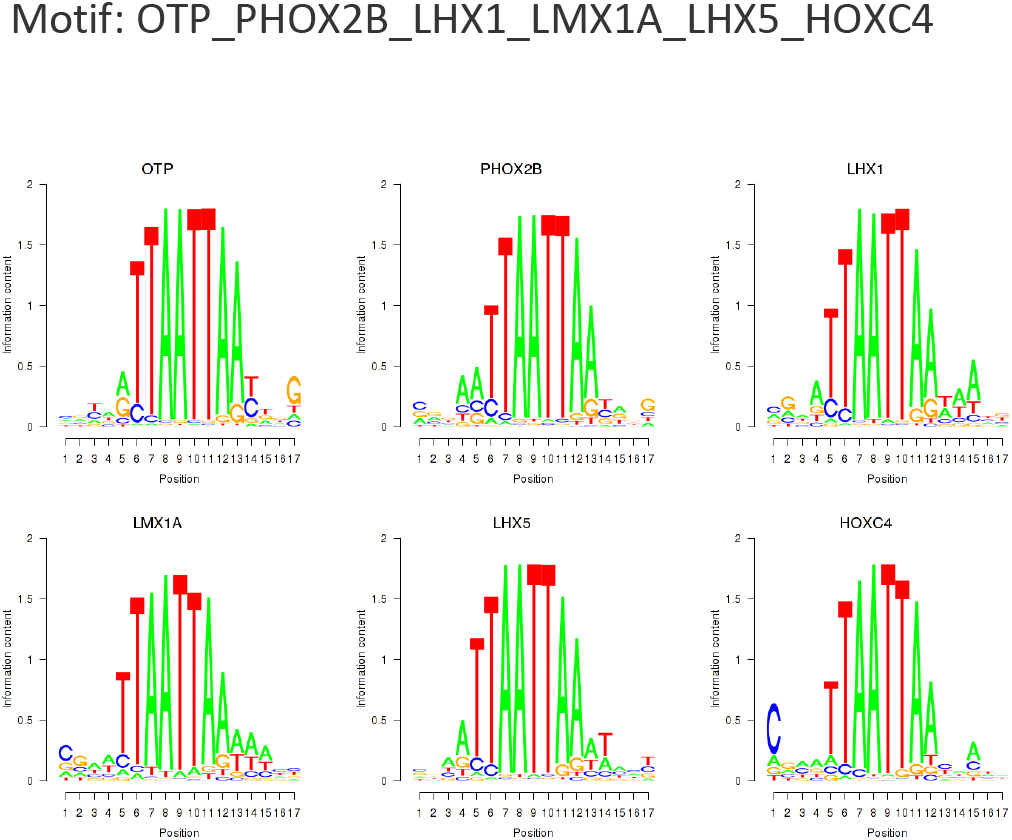
Motif family OTP_PHOX2B_LHX1_LMX1A_LHX5_HOXC4.

**Supplementary Figure 6.**
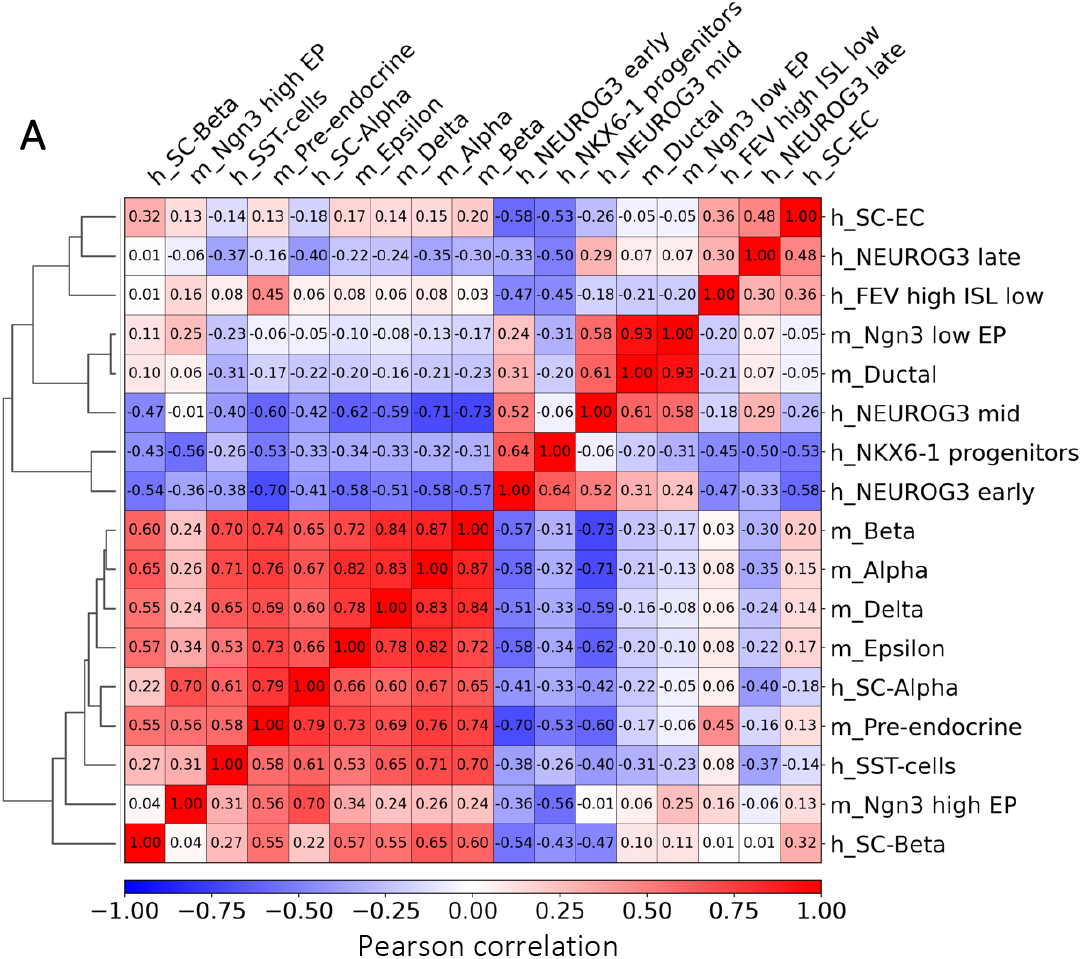
Expression correlation between mouse *in-vivo* and human *in-vitro* datasets. **A**. Matrix plot indicating Pearson correlation among the different clusters from the mouse *in-vivo* dataset (Bastidas-Ponce et al., 2019) and the human *in-vitro* dataset (Veres et al., 2019).

**Supplementary Figure 7.**
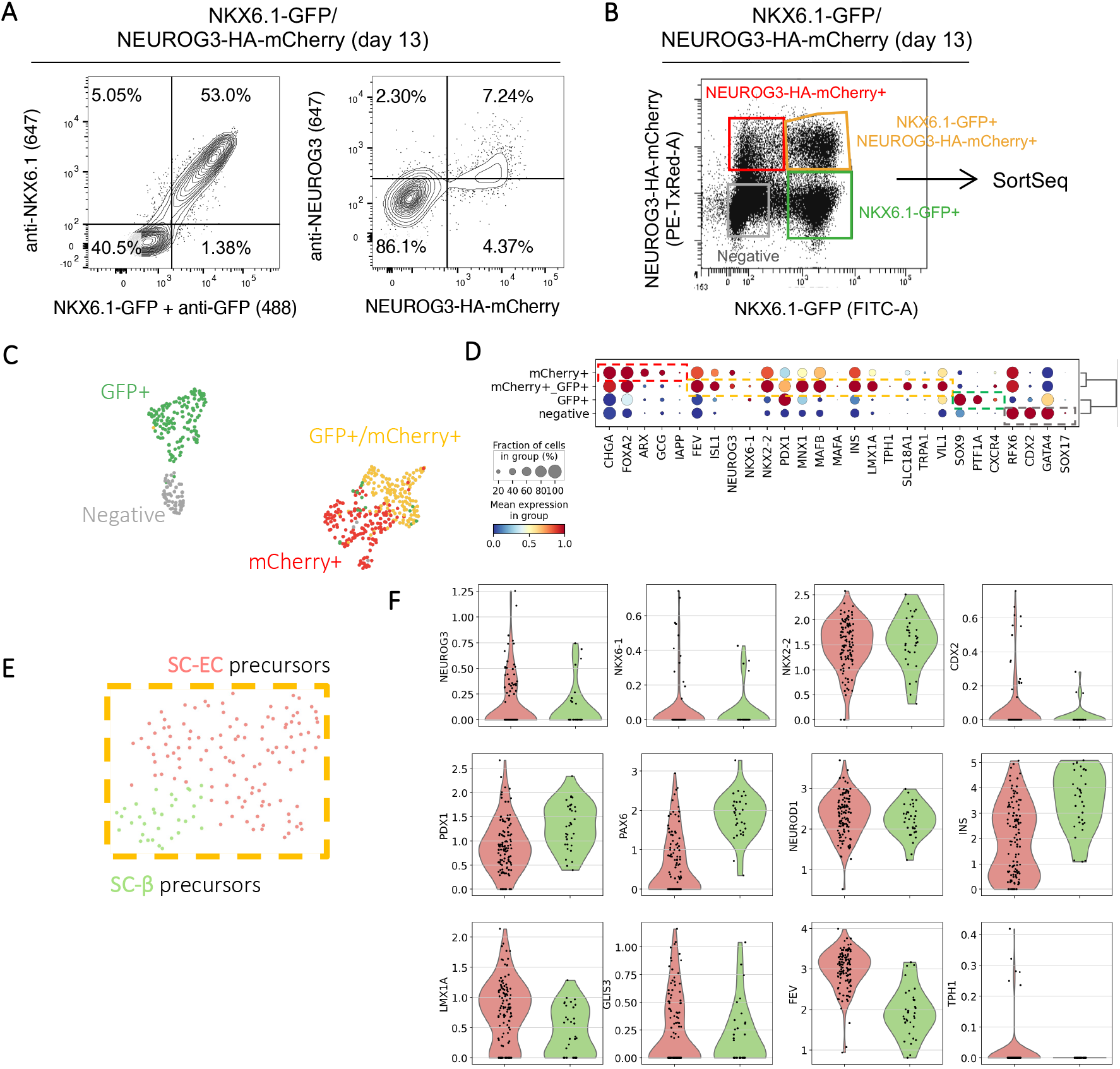
Lineage tracing during *in-vitro* differentiation towards b-like cells. **A**. Representative flow cytometry plots showing the overlap between the GFP or mCherry fluorescent reporters and the NKX6.1 or NEUROG3 proteins they trace, respectively, in NKX6.1- GFP/NEUROG3-HA-mCherry cells at day 13 of differentiation. **B**. Flow cytometry plot showing the gating used to sort living NKX6.1-GFP+ and/or NEUROG3-HA-mCherry+ single cells at day 13 of differentiation for SortSeq single-cell RNA sequencing. **C**. UMAP plot of 522 cells colored according to the fluorescent reporter for which they were sorted. **D**. Dot plot showing expression of known cell-type-specific gene sets. **E**. UMAP plot of 150 cells from the double-positive population. **F**. Violin plots showing expression of known lineage-specific markers in the SC-β precursors vs. SC-EC precursors.

## Methods

### 1. Pipeline

The FateCompass workflow aims to identify key transcription factors (TFs) during a cellular system undergoing differentiation. To mechanistically understand the dynamic transcriptional interactions underlying the cell subtype specification, we reasoned that inherent asynchrony of the cells, coming from single-cell RNA sequencing (scRNAseq) experiments, provides a temporal resolution of the transcriptome; also, that *cis*-regulatory regions of the expressed genes contain essential information of the TFs that regulate their transcription. To this end, we integrated both state-of-the-art methods and newly developed algorithms in a coherent and flexible pipeline. We took as input the gene expression count matrix, X ∈ ℝ^CxG^, where C is the number of cells and G is the number of genes; the velocity field when present, V ∈ ℝ^CxG^; and TFs binding sites predictions in the promoters of the expressed genes, N_gm_ ∈ ℝ^GxM^, where M is the number of TF motifs. Importantly, our pipeline can be generalized to include epigenetic information coming from chromatin accessibility and interactions between promoters and enhancers by extending N_gm_. The FateCompass pipeline consists of three main steps:

i. Retrieve gene expression dynamics of cell differentiation.
ii. Estimate TF activities along the cell-fate decision process.
iii. Identify lineage-specific regulators.

### 1.1 Cell-fate decision dynamics from single-cell RNA sequencing

The main purpose of this work is to study the trajectory a cell follows to arrive to its final state rather than the final state itself. A single cell, whose phenotype is represented by a point in the multidimensional space, will move along a specific trajectory as its composition changes (transcriptomic profile). Considering a regionalized scenario such as the Epigenetic Landscape of Waddington (Waddington, 1957), we reasoned that trajectories converge to end-states which are essentially different from one another; also, that if a cell-system moving along a specific trajectory is pushed slightly out of its way, then the canalization of the landscape will compensate, and eventually, the cell will arrive in the stable state it would typically have done [1]. The process of a cell changing states along a trajectory until it reaches a final fate can be understood as a particle diffusing on a volume. To delineate the differentiation trajectories, we considered two scenarios: one unbiased, in which the diffusing particle, single-cell, follows a random walk under the influence of a vector field, here represented by the RNA velocity until it gets trapped on an attractor; and the other, biased, in which the single-cell is following a random walk from progenitor cells, that are defined as sources, towards mature cells, that are defined as sinks.

#### 1.1.1 Nearest neighbor graph representing the phenotypic manifold

Similar to other methods [2], [3], FateCompass models cell state transitions restricting possible state changes to those consistent with the global structure of the phenotypic manifold via a *k-*nearest neighbor graph based on similarities on the gene expression space. Due to the high sparsity and noise in scRNAseq data, finding nearest neighbors in the raw data using a simple similarity metric is likely to accumulate spurious connections and obscure the structure we seek.

To build the neighbor graph based on solid data trends, we used Uniform Manifold Approximation and Projection (UMAP), a non-linear dimensionality reduction algorithm that estimates the topology of the high dimensional data and uses this information to build a low-dimensional representation that better preserves the local structure of the data over the global variability [4]. Despite the wide use of Principal Component Analysis (PCA) for detecting among-sample heterogeneity, it has shown to be inefficient in dimensionality reduction of scRNAseq data [5], [6], and we reasoned that for large datasets, local and neighborhood structures are more prominent for sample heterogeneity analysis and describe the local structure of the data. Therefore, we choose the non-linear method UMAP, which has proven to be more performant than others like tSNE when embedding in dimensions larger than two [4], which is particularly important when the intention is to use the low dimensional representation for further downstream analysis such as clustering.

Formally, given a dataset, X ∈ ℝ^CxG^, a *k*-nearest neighbor graph is constructed using the Euclidean distance on the D dimensions of the UMAP embedding, where 2 < D <K, and K is the number of neighbors initially used for the embedding. The reason for this is that UMAP will have significantly diminishing returns as it approaches the value of *n_neighbors* used for the embedding [4]. Then, to build the adjacency matrix, M, we kept the same number of neighbors for each cell, NN; and the distances were weighted equally since the intermediary embedding step favored a single strong connection vs. lots of weak links.

#### 1.1.2 Modeling transition probabilities using a Markov process

Single-cell transcriptomics provides a static picture of a time-evolving system whose possible states are represented by points in a manifold, in other words, X(t) ≡ state point of the system at time t. The value of X at some initial time t_0_ is fixed, X(t_0_) = x_0_, and for successive instants t_1_, t_2_, …, t_n_; where t <t_2_< … <t_n_ ; there are n corresponding random states X(t_1_), X(t_2_), ., X(t_n_). To determine how the cells move from progenitors to mature cells, we assumed that they traverse the manifold in small steps under the influence of an external force, drift, in the direction of differentiation, e.g., RNA velocity or gradient of potential energy from progenitor cells (sources) to mature cells (sinks). This can be modeled using a Markov chain to represent cell fate choices in a probabilistic manner as follows [7]:

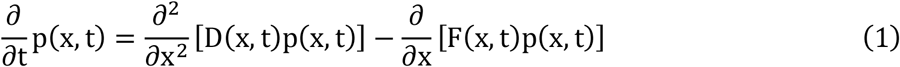

where the left-hand-side of the equation represents the probability of being at the state x at the time t, p(x, t); the first term on the right-hand-side is the flux through the state x due to diffusion, D(x, t), and the second term on the right-hand-side is the flux through the state due the external force or drift, F(x, t).

To outline the differentiation trajectories, we took advantage of the system’s stability in the observable space. We went from the continuous-space previously described (equation 1) to a discrete space with *state-dependent* drift criteria. We decided to move to a discrete space due to the ill-posed nature of the problem and that the observed states are not enough to constrain the solution. As a result, the gene expression dynamics are described by a discrete Markov process on a network. This simplification avoids the complex problem of inferring the high-dimensional drift field, F(x, t), with only a few thousand observations. In this way, we allowed jumps only to the observed states, with state-dependent drift, F(x_i_), and the weight of each jump given by the normalized transition probability, ∏(x_j_|x_i_, F(x_i_)). Below we describe the form of the propagator when considering different drifts, namely, RNA velocity and Potential energy.

#### 1.1.3 RNA velocity as driving force

To get the transition probabilities using RNA velocity information, we reasoned from equation (1) that the drift directing the state-transition probabilities is given by the direction of differentiation, represented by the direction of the velocity vector [8]. In addition, we made the following assumptions: for a very small dt, we approximate dt → Δt, we assumed the diffusion coefficient to be completely homogeneous (D), and the drift to be time-independent and locally constant around a given state x (V(x)). Under these assumptions, and in the continuous space, the propagator of equation (1) can be approximated by Gaussian distribution:

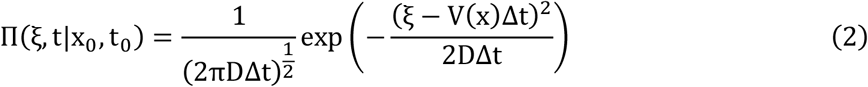

where ξ is the distance between the current state and the next possible state, ξ= |x_t_ − x_0_|. A row normalization is applied to transform the Gaussian distribution into transition probabilities over the network of discrete observed states:

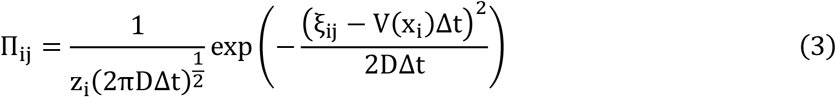

with row normalization factors z_i_ = ∑_j_ Π_*ij*_.

We fitted the diffusion coefficient, D, and Δt heuristically based on the number of neighbors. Shortly, we set Δt such that on average the number of nearest neighbors can be reached, and D such that the average number of connections is twice the number of nearest neighbors. In this way, the distance traveled until the next state is close to the distance to the nearest neighbors, and the explored neighborhood is within the velocity gradient. Therefore, we make sure that the progression of stable-states follows the direction of differentiation.

#### 1.1.4 Potential energy as driving force

To estimate transition probabilities using the gradient of potential energy from progenitor cells (sources) to mature cells (sinks) we defined the boundaries of the phenotypic space, i.e., we fixed the starting point (source) and the endpoints (sinks). In this case, we built the energy landscape by doing a force balance between each *i* point in the network (observable state) and the k states of interest (sources and sinks). Briefly, we assumed the force at each point on the network, 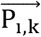, to be directly proportional to an attraction or repulsion coefficient, Q_k_, and inversely proportional to the distance of that given state to the state of interest, dist(x_i_, x_k_). This is equivalent to:

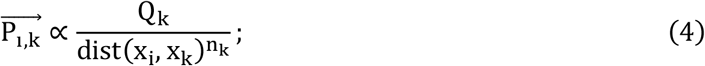

with i ∈ [1, C] and k := sources & sinks. Then, assuming the energy landscape to be time-independent and locally constant around a given state i, the force balance around each i gives the energy landscape, W, as follows:

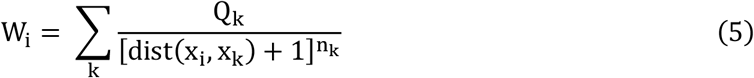

To approximate this, we assume |Q_sources_| = |Q_s*i*nks_| = |Q| and _sources_ = _sinks_ =n, and we used a heuristic criterion to set their value based on the distribution profile of the energy landscape. We reasoned that the potential energy gradient should be high enough for the sources to be phenotypically distinct from the sinks. Having the potential energy for each state, we defined the transition kernel as [9]:

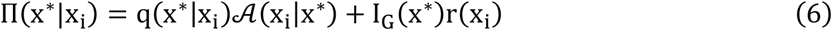

where the left-hand-side of the equation, n(x*|x_i_), is the transition probability from x_i_ to x*, the first term on the right-hand-side represents the probability of jumping from state x_i_ to state x given by the multiplication of the proposal distribution q and the acceptance distribution 𝒜, and the second term on the right-hand-side is the probability of not jumping represented by the rejection distribution r. Having a symmetric proposal distribution,q (x* | x_i_) = (x_i_| x*) = 1/NN, with NN the number of nearest-neighbors, the acceptance probability is

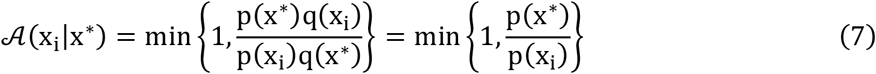

where p is the invariant distribution,

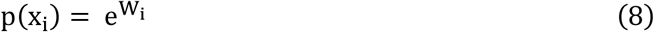

Finally, the rejection distribution reads

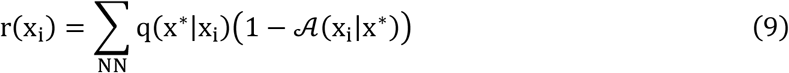

#### 1.1.5 Stochastic simulations

To describe the time evolution of the previously described Markov processes, we used a numerical approach called Monte Carlo sampling algorithm. Shortly, the idea of a Monte Carlo simulation is to

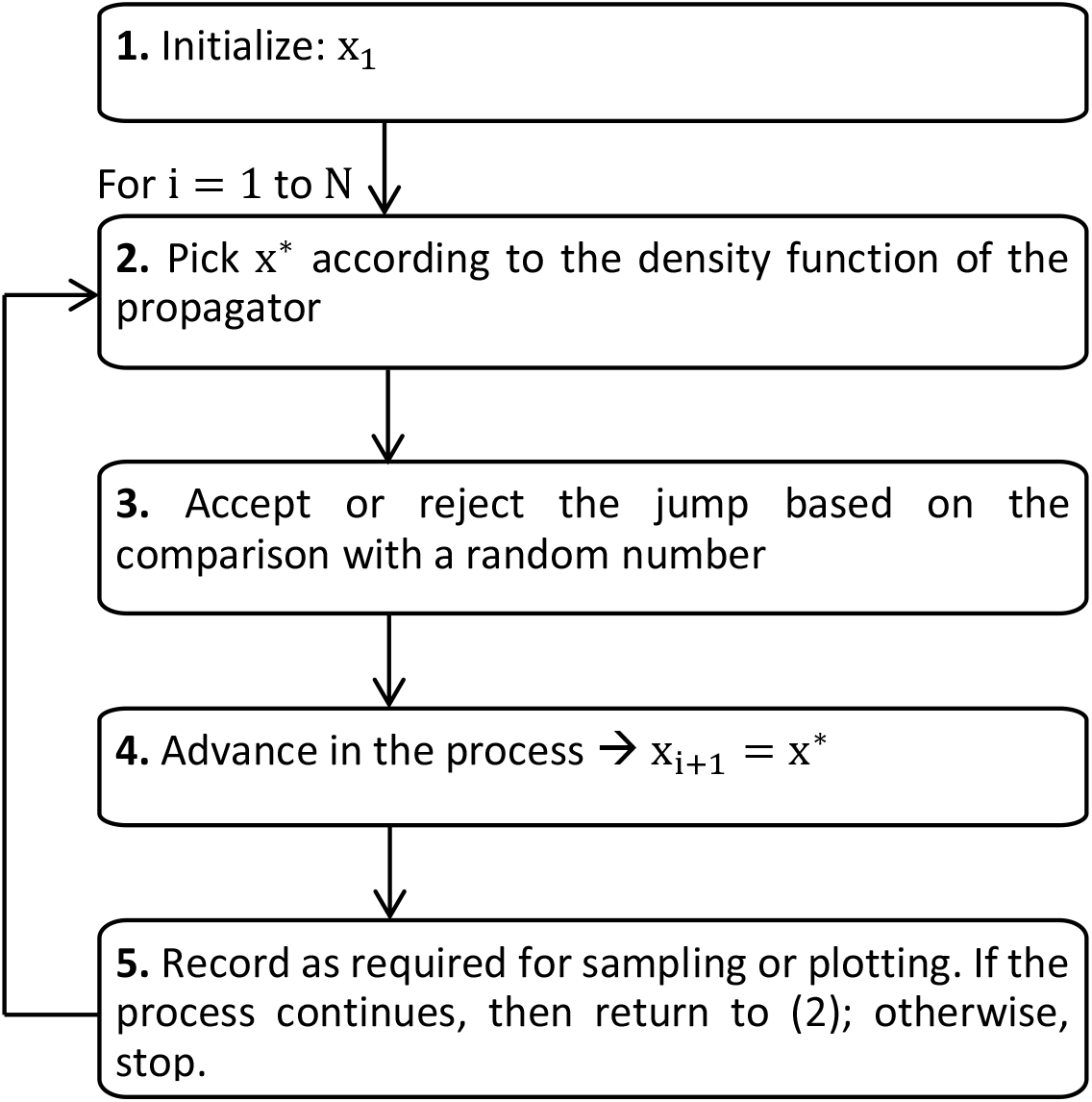

draw an i.i.d. set of samples 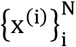 from a target density p(x) [9]. To estimate the values of X withoutknowing the density function p(x), we used sampling methods that essentially mimic the real-time evolution of the process. The pseudo-code for the simulations is below:

#### 1.1.6 Average profiles over stochastic trajectories

We used the previously generated N samples with the following empirical point-mass function to approximate the expected value of the final quantities of interest, mean and standard deviation, as follows,

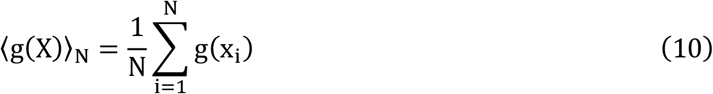

and

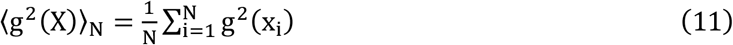

Of note, the estimates (10) and (11) will become exact in the limit N → ∞.

#### 1.1.7 Fate probabilities

We defined the fate probabilities based on the information of the stochastic trajectories. We reasoned that if one of the simulated random walks on the Markov chain passes by cell i; then, we continue to simulate the random walk until it reaches any cell from a final sink, and we record this for every single cell on the graph. In the end, we estimated the fate probabilities by counting how often a random walk that visits cell i terminated in any of the terminal index sinks.

### 1.2 Modeling regulatory interactions between transcription factors and *cis*-regulatory regions

To model the regulatory interactions underlying cell-fate decisions, we considered TFs as the central drivers of transcriptional regulation. TFs are usually designed to transit rapidly between active and inactive molecular states at a rate modulated by a specific environmental signal. Each active TF can bind the DNA to regulate the rate at which specific target genes are transcribed [10]. This section describes the model we used to infer TF activates in single cells from their gene expression profiles.

#### 1.2.1 Binding site matrix

First, to find the regulatory interactions between TFs and the different genes, we used TF binding sites predictions reported in [11], and available in the Swiss Regulon Portal (https://swissregulon.unibas.ch/sr/downloads). Briefly, it uses a Bayesian framework to estimate the posterior probability that a binding site for a given weight matrix (associated with a motif) occurs in an interval. Next, we summarized the TF binding sites in a matrix of site-counts by summing the posterior probabilities for each motif in the promoter of each gene. We defined a promoter as the TSS +/- 1kb, Fig. 1d.

#### 1.2.2 Linear model to estimate motif activities

We hypothesize that the expression level of each gene is proportional to the activity of the TFs that can potentially bind to its promoter. Therefore, as in the original framework [12], we modeled the log-expression level of a gene as the linear combination of motif activities weighted by their number of the binding sites present in its promoter, that is,

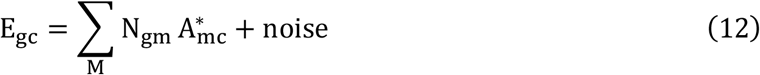

where E_gc_ are the cell- and gene-normalized log-expression values, N_gm_ are the motif-normalized site-counts, and 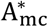 are the cell-normalized motif activities. The term noise is related to multiple sources, namely technical, biological, and error in the model. To estimate the unknown motif activities 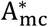, we first used minimum norm least-squares (*lsqminnorm*) solution to linear equations implemented in Matlab 2018b to fit the best estimates of 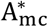 from equation 12. Briefly, *lsqminnorm* uses the complete orthogonal decomposition to find a low-rank approximation of N_gm_. Next, to control the model’s complexity and avoid overfitting, we applied regularization as explained in the following section.

#### 1.2.3 Regularization using data diffusion

A model’s ability to reproduce intricate patterns in data is typically related to its number of parameters and complexity. However, the higher the complexity of a model, the higher the risk of overfitting, i.e., fitting spurious noise in the data leading to poor generalizing performance when applied to new observations. Single-cell transcriptomic data present large technical and biological noise introducing variability in the gene expression profiles across cells that do not reflect true variability in the physiological cellular state. Not accounted for, this variability propagates to the inferred TF activities leading to non-functional cell-to-cell differences in TF activates levels. To control for this, we penalized the model’s complexity by introducing a regularization term that enforces the smoothness and stability of the fitted activities across cells. We embedded the cells in a low-dimensional manifold that faithfully represents the phenotypic similarities (section 1.1.1). Next, we imputed a cell’s motif activities as the weighted average of the activities across the neighboring cells. This strategy, which is mathematically akin to diffusing heat through the data, has been used to correct for dropout and other noise sources on transcriptomic data [13]. It reads,

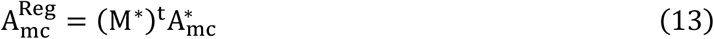

where 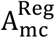 are the regularized activities, M* is the transition matrix, and 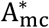 are the maximum-likelihood estimates of the activities. Raising M* to the power of t results in a matrix where each entry represents the probability that a random walk of length t starting at cell i will reach cell, a process similar to diffusion [13]. Importantly, we want a cell’s own observed values to have the highest impact on the imputation of its own values; therefore, our transition matrix M* allows for self-loops, and these are the most probable steps in the random walk. Thus,

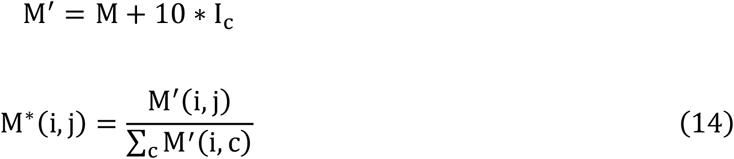

To find the optimal t, we evaluated the impact of t on the final imputed data. We used an 80/20 cross-validation scheme, where the set of promoters was divided randomly into two sets, the training set containing 80% of all promoters and the test set with the remaining 20%. We used the training set to fit the motif activities and the test set to evaluate the quality of the fit. Then, we choose the value of t that minimizes the mean square error (MSE) between the observed expression levels and those predicted by the model in the test set.

### 1.3 Differential motif activity analysis

To identify key lineage-specific regulators during the cell-fate decision process, we defined a differential motif activity analysis based on the following criteria: **(i)** motifs with the high positive z-score, i.e., motifs that significantly varied across cells compared with their estimated errors. **(ii)** motifs with high activity variability across the linage-specific differentiation trajectory. Finally, **(iii)** motifs with a high temporal correlation between its activity and mRNA expression within a specific window of time lags.

#### 1.3.1 Z-score

To estimate the importance of a motif, we reasoned that activities that fluctuate the most across conditions should be the more important. Therefore, we used the number of standard-deviations that the activity of motif m is away from its average of zero corrected by the precision of the estimation (error bar), also known as z-score, as an indicator of the importance of each motif

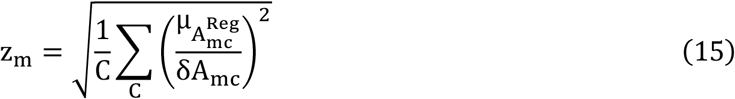

where C is the number of cells, 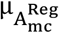 is the mean of the motif activity distribution, and δ A_mc_ is the reliability of the fitting of 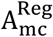 (error-bar) [12]. Since we do not know the posterior distribution of the 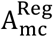, there is no analytical way to estimate the standard deviations δA_mc_. Therefore, we used bootstrapping, as explained below.

##### Bootstrapping

Important to the concept of bootstrapping is that inference about a population from sample data can be modeled by resampling the sample data and performing inference about a sample from the resampled data. Then, we built the distribution of the estimate for the motif activities using random sampling with replacement following the steps below:

i. The activity of motif in cell c is a function of the gene expression on that cell and the binding-site matrix; in other words,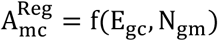. Similar to the cross-validation scheme, here we sampled taking the promoters/genes as observations.
ii. Resample from E_gc_ and N_gm_ taking randomly 80% of the observations. Importantly, the bootstrap resample has the same number of observations as the original data used in the training (sections 1.2.2 & 1.2.3).
iii. Compute the estimate of 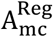 .
iv. Repeat (ii) and (iii) a large number of times, B, to get 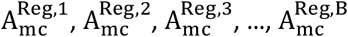.
v. Use the estimates in (iv) to build the empirical bootstrap distribution of the estimate for the motif activities.
vi. Infer 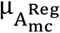 and δA_mc_ from the empirical bootstrap distribution of the estimate.

#### 1.3.2 Variability over time

We seek TFs with a high rate of change in the activity with respect to time, which is, clearly, the definition of the first-order derivative. However, since we do not know the exact distribution of the motif activities, there is no easy way to get the analytical solution. To get a proxy of the rate of change, we estimate the standard deviation over time of the mean activity profile along the simulated trajectories (computed via Eqn 10), as follows:

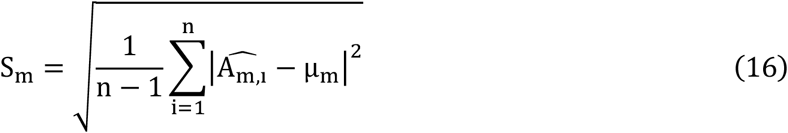

where n is the number of iterations, 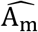 is the average activity of motif m over the simulated trajectories, and µ_m_ is the mean of 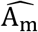

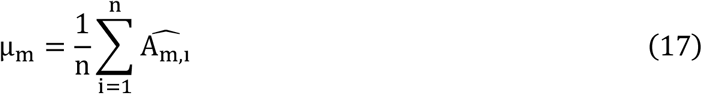

#### 1.3.3 Dynamic correlation

Next, we used cross-correlation to identify dynamical correlations between average motif activities and their average mRNA expression along the differentiation trajectories. *Cross-correlation* is defined as a similarity measure between two series as a function of the displacement of one relative to the other.

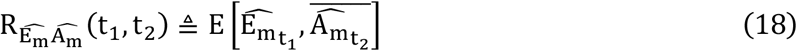

the left-hand-side reads as the cross-correlation between times t_1_ and t_2_ for the average mRNA expression over the simulated trajectories of, 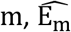, and 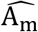. Next, we converted cross-correlation to Pearson correlation to facilitate the comparisons (1: maximum correlation, 0: no correlation, and -1: maximum anti-correlation).

### 2. Mouse in-vivo dataset from endocrinogenesis

We used scRNAseq data from the developing pancreas during the secondary transition, i.e., from embryonic day 12.5 to 15.5, published by Bastidas-Ponce et al., 2019 [14] and available in the Gene Expression Omnibus under accession number GSE132188. In particular, we used data from the last experimental time point, E15.5. We retrieve an annotated object directly from https://scvelo.org with the expression matrix and the annotations for unspliced/spliced reads using the following command: scvelo.datasets.pancreatic_endocrinogenesis(). Our final subset for Figure 2 contained 3696 cells. We kept the original cluster annotation reported by Bastidas-Ponce et al., 2019.

#### Data pre-processing and velocity computation

We used SCANPY [15] and scVelo [16] with most of the default parameters. We filtered out genes with less than 20 counts in both spliced and unspliced layers. Next, we normalized by total counts per cell, log-transformed the data, and kept the top 2000 highly variable genes. We embedded the data in the PCA space and used the top 30 principal components to compute a k-nearest neighbor graph with k = For visualization, we used UMAP embedding with two dimensions with default parameters. To compute RNA velocity, we used scVelo’s dynamic model of splicing kinetics.

#### FateCompass specific computations

For downstream analysis of the FateCompass pipeline, we embedded the mouse in-vivo data of gene expression and RNA velocity on ten dimensions in the UMAP space. Next, we computed a neighborhood graph in the reduced gene expression space with k = 10. This setting was the graph structure for the Markov chain operations of the FateCompass pipeline. The edges of the Markov chain were directed using the RNA velocity information and equation 3. We outlined stochastic trajectories using a Gibbs Sampling algorithm. Last, the thresholds for the differential motif activity analysis were: minimum z-score of 1.5, minimum standard deviation over trajectories of 0.003, and minimum Pearson correlation of 0.7.

### 3. Human *in-vitro* dataset from differentiation towards β-like cells

We used a scRNAseq time-series dataset from a differentiation protocol from human embryonic stem cells towards β-like cells profiled using inDrops [17]. The differentiation protocol consists of six stages, with pancreatic endocrine cells appearing throughout stage five. Veres et al. (2019) performed sequencing at the end of each stage and daily sampling across stage five. The data is available in the Gene Expression Omnibus under accession number GSE114412. We restricted the data to the endocrine lineage, from NKX6-1+ progenitors to hormone-producing cells. Our final subset for figure 3 contained 25299 cells. We kept the original cluster annotations.

#### Data pre-processing and velocity computation

Sequencing reads were preprocessed according to the dropEst pipeline (https://github.com/kharchenkolab/dropEst). A reference index was built from the Ensembl GRCh38 human genome assembly and the GRCh38.88 transcriptome annotation to run the pipeline. Shortly, we first extracted the cell barcodes and UMIs from the library using the dropTag command. Next, we used STAR 2.7.9a to map the reads to the human transcriptome. Finally, we used the dropEst command with the option -V, which allows the output of three separate matrices containing only UMIs of a specific type: intronic, exonic, or exon/intron spanning. These matrices were used to build an annotated h5ad object with the unspliced layer equal to the sum of intronic and spanning UMIs and the spliced layer corresponding to the exonic UMIs. We used SCANPY and scVelo with mostly default parameters. We filtered genes to be expressed in at least three cells. Next, we normalized by total counts per cell, log-transformed the data, and kept the top 2000 highly variable genes. We embedded the data in the PCA space and used the top 50 principal components to compute a k-nearest neighbor graph with k = 50. For visualization, we used UMAP embedding with two dimensions with default parameters. To compute RNA velocity, we used scVelo’s dynamic model of splicing kinetics.

#### FateCompass specific computations

For downstream analysis of the FateCompass pipeline, we embedded the human in-vitro data of gene expression and RNA velocity on ten dimensions in the UMAP space. Next, we computed a neighborhood graph in the reduced gene expression space with k = 50. This setting was the graph structure for the Markov chain operations of the FateCompass pipeline. The edges of the Markov chain were directed using the potential energy landscape described in equation 5. We outlined stochastic trajectories using a Monte Carlo Sampling algorithm. Last, the thresholds for the differential motif activity analysis were: minimum z-score of 1.5, minimum standard deviation over trajectories of 0.006, and minimum Pearson correlation of 0.7.

## 4. Lineage tracing of NEUROG3+/NKX6-1+ cells

### Generation of the hiPSC line, differentiation to pancreatic endocrine progenitors

The NKX6.1-GFP iPSC line (1a-21) in which the NKX6.1 coding region is fused to a T2A-eGFP reporter is described in Gupta et al (2018) and was obtained through the IMI/EU sponsored StemBANCC consortium via Christian Honoré (Novo Nordisk A/S) [18]. This line was gene edited by CRISPR/Cas9 to knock in a 3xHA-P2A-3xNLS-mCherry cassette in fusion with NEUROG3 coding sequence, following exactly the strategy detailed in Schreiber et al (2021) [19]. Picked clones were characterized by PCR genotyping, sequencing and ddPCR. The heterozygous NKX6.1-GFP/NEUROG3-HA-mCherry clone#10 was differentiated to pancreatic endocrine progenitors (Stage 5 day3, or day 13) following the protocol detailed in Schreiber et al (2021), based on Petersen et al (2017) [19], [20]. Flow cytometry analyses were performed on differentiated cells as described in Schreiber et al (2021) using sheep anti-NEUROG3 (R&D), chicken anti-GFP (Abcam), Alexa-647 coupled anti-NKX6.1 (BD), Alexa-647 Donkey anti Sheep (Jackson ImmunoRes) and Alexa-488 Donkey anti Chicken (Jackson ImmunoRes) antibodies.

### FACS sorting and single-cell RNA sequencing

Cells were harvested with TrypLE Select and sorted using a FACSAria Fusion cell sorter (BD) directly into 384-well plates with ERCC spike-ins (Agilent), reverse transcription primers and dNTPs (both Promega). Single cell sequencing was performed according to the Sort-seq method [21] by Single Cell Discoveries. Briefly, Sequencing libraries were generated with TruSeq small RNA primers (Illumina) and sequenced paired-end at 60 and 26 bp read length, respectively, on the Illumina NextSeq. Reads were mapped to the human GRCh38 genome assembly. Sort-seq read counts were filtered to exclude reads with identical library-, cell- and molecule barcodes. UMI counts were adjusted using Poisson counting statistics [21].

### Data pre-processing

We used SCANPY with mostly default parameters. We removed cells with a high fraction of mitochondrial gene counts (>50%), and a high percentage of ERCC spike-in reads (>50%); also, cells with more than 100000 and less than 10000 counts were excluded. We filtered genes to be expressed in at least three cells and remove spike-in genes for downstream analysis. Next, we normalized by total counts per cell, log-transformed the data. We embedded the data in the PCA space and used the top 15 principal components to compute a k-nearest neighbor graph with k = 15. For visualization, we used UMAP embedding with two dimensions with default parameters. We used Louvain-based clustering [22] for clustering and cell-type identification. Cell types were annotated based on the expression of known marker genes.

### Motif activity estimation

Motif activities were estimated according to the procedure described in section 1.2. We embedded the data of gene expression on ten dimensions in the UMAP space. Next, we computed a neighborhood graph in the reduced gene expression space with k = 10. This setting was the graph structure for the regularization of the motif activities.

